# Sarand: Exploring Antimicrobial Resistance Gene Neighborhoods in Complex Metagenomic Assembly Graphs

**DOI:** 10.1101/2023.10.29.564611

**Authors:** Somayeh Kafaie, Robert G. Beiko, Finlay Maguire

**Affiliations:** Faculty of Computer Science, Dalhousie University, 6050 University Avenue, Halifax, Nova Scotia B3H 4R2, Canada; Department of Mathematics and Computer Science, Saint Mary’s University, Halifax, Nova Scotia, Canada; Department of Community Health and Epidemiology, Dalhousie University, 6050 University Avenue, Halifax, Nova Scotia B3H 4R2, Canada; Institute for Comparative Genomics, Dalhousie University, 6050 University Avenue, Halifax, Nova Scotia B3H 4R2, Canada

## Abstract

Antimicrobial resistance (AMR) is a major global challenge to human and animal health. The genomic element (e.g., chromosome, plasmid, and genomic islands) and neighbouring genes associated with an AMR gene play a major role in its function, regulation, evolution, and propensity to undergo lateral gene transfer. Therefore, characterising these genomic contexts is vital to effective AMR surveillance, risk assessment, and stewardship. Metagenomic sequencing is widely used to identify AMR genes in microbial communities, but analysis of short-read data offers fragmentary information that lacks this critical contextual information. Alternatively, metagenomic assembly, in which a complex assembly graph is generated and condensed into contigs, provides some contextual information but systematically fails to recover many mobile genetic elements. Here we introduce Sarand, a method that combines the sensitivity of read-based methods with the genomic context offered by assemblies by extracting AMR genes and their associated context directly from metagenomic assembly graphs. Sarand combines BLAST-based homology searches with coverage statistics to sensitively identify and visualise AMR gene contexts while minimising inference of chimeric contexts. Using both real and simulated metagenomic data, we show that Sarand outperforms metagenomic assembly and recently developed graph-based tools in terms of precision and sensitivity for this problem. Sarand (https://github.com/beiko-lab/sarand) enables effective extraction of metagenomic AMR gene contexts to better characterize AMR evolutionary dynamics within complex microbial communities.

## INTRODUCTION

Antimicrobial resistance (AMR) is a major global health threat (1) with an estimated 1.27 million deaths attributed to AMR in 2019 alone (2). Lateral gene transfer (LGT) of mobile genetic elements (MGEs), such as plasmids and genomic islands, can drive the spread of AMR between organisms and habitats (3). For example, the dynamics of colistin resistance amongst livestock-associated microbial communities influences the dynamics of colistin resistance in clinically-associated pathogens (4). To develop effective interventions to control AMR in this inter-sectoral “One Health” context, we need to be able to characterise the evolution and transmission of AMR within and between microbial communities. An AMR gene’s associated genomic element(s) (e.g., chromosome, plasmid, genomic island) and neighbouring genes can play key roles in its function (5), regulation (6, 7), evolution (8), and likelihood of undergoing LGT (9, 10). Therefore, to mitigate AMR we require tools which can systematically explore the genomic context (or neighbourhood) of AMR genes in genomes within key microbial ecosystems such as human and animal microbiomes, agricultural soil, and wastewater.

The isolation and sequencing of individual genomes from these communities can provide an inventory of AMR genes and their contexts. However, even with advances in single-cell methods (11) and culturing approaches (12, 13) this is only feasible for a subset of the microbes in most host-associated or environmental communities. Alternatively, metagenomics, in which all the DNA present in a community is simultaneously sampled and sequenced, can be used to characterise complex microbial communities. Although long-read sequencing technologies such as nanopore (14) and single-molecule real-time sequencing (15) are now used for metagenomic sequencing (16, 17), the greater throughput of relatively short-read technologies (150-250bp) means they still comprise the majority of metagenomic data sets. As these reads are shorter than most genes, it is only rarely possible to identify specific AMR alleles or recover genomic context information directly from these reads.

Metagenomic assembly, in which shared sequences between reads are used to recover longer sequences, offers one possible solution to this problem. Assembly algorithms aim to construct graphs that represent candidate orderings of sequence reads (or their constituent k-mers), then condense these graphs to produce unambiguous contiguous sequences (*contigs*). Sequencing errors, repetitive DNA segments, and other structural variants make even isolate genome assembly a challenging task (18), and the MGEs that often bear AMR genes are among the most difficult parts of genomes to correctly assemble (19). Metagenomic assembly is even more complicated, with the uneven abundance of organisms in a sample, genome differences at the strain level, and nearly identical genome segments across closely related species leading to substantial uncertainty and ambiguity during assembly (18). This often leads to metagenomic assembly recovering missing, partial, or incorrect gene neighborhoods around target genes (19, 20).

Direct exploration of the sequence graph generated during metagenomic assembly (before it is condensed into linear contigs) offers an alternative and potentially more sensitive approach to identify and explore the genomic context of key genes within microbial communities (21). The promise of this approach has been shown in related work resolving closely related strains (22), SNP calling (23), and performing rapid gene homology searches (24) in complex metagenomes.

Tools for graph-based genomic context analyses can take a localised assembly approach (such as in MetaCherchant) in which reads and k-mers related to the query gene(s) are identified and then a local de Bruijn graph constructed representing the query neighborhood (25). However, while effective for capturing the metagenomic diversity of the query genes themselves, these localised assembly approaches may be limited in their ability to capture the wider gene neighbourhood(s). Alternatively, other methods first construct the entire assembly graph and then extract the query neighborhood either manually through scaffolding (21) or through automated graph-theoretic approaches to subgraph extraction (such as in Spacegraphcats) (26, 27). However, these extracted subgraphs contain many possible upstream and downstream paths and lack an easy approach to resolve the true genomic neighbourhoods (i.e., correctly join upstream and downstream paths) from false chimeric path pairs that do not exist in any underlying genomes. This problem is particularly acute for mobile AMR genes as they may have multiple genomic contexts within a metagenome, and be associated with repetitive sequences that are among the most difficult-to-assemble parts of bacterial genomes.

To try and address these challenges, we developed Sarand (https://github.com/beiko-lab/sarand), a new method to extract the genomic neighborhoods of AMR genes from metagenomic assembly graphs. Sarand uses BLAST-based homology searches to identify AMR gene-associated nodes in the assembly graph, enumerates local upstream and downstream paths, and then applies coverage-based statistics to filter out likely chimeric neighbourhoods. To enhance exploration and comparison of AMR genes within sequenced microbial communities, Sarand also annotates and visualises the extracted gene neighbourhoods. We evaluated

Sarand’s ability to reconstruct AMR gene neighbourhoods compared to metagenomic assembly, local assembly-based MetaCherchant (25), and the subgraph-based Spacegraphcats (26) using low, medium and high diversity simulated metagenomes as well as 3 real urban sewage metagenomes from Hendriksen et. al., (28). These analyses show that Sarand had superior sensitivity (47% and 134% average improvement over assembled contigs and MetaCherchant respectively) in identifying AMR gene neighbourhoods while also outperforming other approaches in avoiding chimeric/false neighbourhoods (46% and 48% respective improvement).

## MATERIALS AND METHODS

### Overview of Sarand

As shown in Figure 1, Sarand starts with a coverage-annotated metagenomic assembly graph in GFA format generated by tools such as metaSPAdes (36), BCALM (27), or megahit (37). In these assembly graphs, fragments of DNA sequences are presented as nodes (i.e., segments), while overlapping segments are connected via edges. A reference set of curated AMR genes is downloaded from the Comprehensive Antibiotic Resistance Database (CARD) (30), and aligned against the assembly graph using the graph-based BLASTN implementation in Bandage (31) (or optionally via an alternative GraphAligner query (32)). Then, for each predicted AMR gene, all paths of the graph within a given sequence length upstream and downstream of the nodes representing the AMR gene are traversed and extracted. These upstream and downstream paths are then annotated using either Bakta (33) or Prokka (34) and RGI (38). To generate genomic neighbourhoods, upstream and downstream paths are paired into valid neighbourhoods using a differential gene coverage statistic. Finally, the dna-features-viewer (35) library is used to visualize the annotation of the extracted and filtered genomic neighbourhoods.

**Figure 1.**
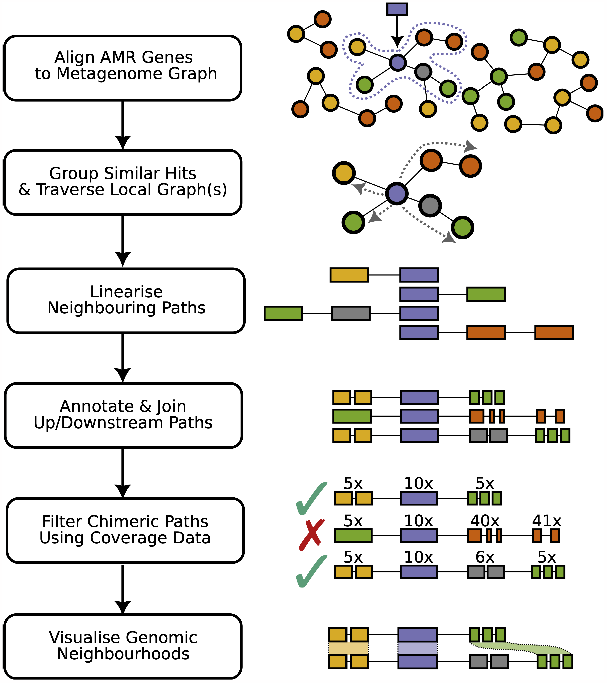
Overview of Sarand. Sarand ingests a GFA-formatted metagenome assembly graph with node coverage annotation. BLASTN-based (29) homology search of the graph is then performed using nucleotide “homology model” sequences from the Comprehensive Antibiotic Resistance Database (CARD) (30) and optionally either Bandage’s (31) graph-based BLAST implementation or GraphAligner (32). Similar hits are grouped using sequence identity. The graph is then traversed for each group to extract the linear upstream and downstream neighbouring paths of the predicted AMR genes (within a user-specified sequence length). These upstream and downstream paths are annotated using Bakta (33) or Prokka (34) and CARD’s RGI (30) before being paired into neighbourhoods. False neighbourhoods that do not exist in the underlying genomes (i.e., incorrectly matched upstream and downstream paths) are then removed using a differential gene coverage statistic. Finally, extracted AMR neighbourhoods are visualised using the dna-features-viewer library (35).

### AMR alignment and grouping

The Bandage implementation of BLASTN returns a list of all possible paths in the graph aligning to reference AMR gene sequences with their corresponding sequence identity and query coverage. We retain all paths with identity and coverage above a given user-supplied threshold (the default value for both parameters in Sarand’s setting is 95%). Similar AMR sequences (e.g., alleles of New Delhi metallo-*β*-lactamase which differ by single amino acids) may align to the same graph nodes. To avoid redundancy, we group these hits together on the basis of sharing > 95% of the same graph paths and perform the neighborhood sequence extraction task only once for each group.

### Extraction of gene neighborhoods

To extract the neighborhood sequences of an AMR gene in an assembly graph, a depth-first recursive search is used to extract all sequences of upstream and downstream nodes up to a predefined sequence length (e.g., 1000bp). For upstream extraction, we start from the first node matching the AMR sequence in the graph. We recursively add upstream nodes until the extracted path reaches a dead-end node (i.e., an upstream node with no incoming edge or downstream node with no outgoing edge) or its sequence is extracted up to the user-defined neighbourhood size. Similarly, the downstream sequences are extracted starting from the last node in the AMR path. We allow users to optionally set a threshold time (e.g, 1 hour) for enumeration of all upstream and downstream paths; if this threshold is exceeded then we terminate the search and return only those paths that were identified thus far.

Our graph traversal rules are defined based on node orientation. The orientation of each node represents the direction of its sequence, where ‘-’ and ‘+’ refer to negative and positive orientation respectively. The sequence of a node with ‘-’ orientation is defined as the reverse complement of its original sequence. Two edges (e.g., *A* → *B* and *B* → *C*) can be connected directly in a path (e.g., *A* → *B* → *C*) only if the orientation of the common node (i.e., *B*) is the same in both edges. For example, with edges *A*^+^→ *B*^−^and *B*^+^→ *C*^+^, there is no path between *A* and *C* through *B*, because of inconsistency in the orientation of node *B*. However, if we have edge *B*−→ *C*^+^as well, path *A*^+^,*B*^−^,*C*^+^ can be extracted. This is a key rule that needs to be followed during neighborhood extraction, and turns some nodes to dead-ends. The edge *A*^−^→ *B*^+^→, for example, can be reversed and converted to edge *B*^−^ → *A*^+^ in an assembly graph if it can help to extend a path. The only difference is that the latter edge represents the reverse complement sequence of the former edge. Therefore, the main rules for extracting upstream (downstream) paths of a given node, *B*, are as follows:

- If there is an edge *C*→*B* (*B*→*C*) where the orientation of *B* in this edge is the same as its orientation in the part of the path that has already been extracted, *C* is eligible to be considered in the upstream (downstream) path.
- If there is an edge *B*→*C* (*C*→*B*) where the orientation of *B* in this edge is the reverse of its orientation in the already extracted part of the path, *C* with the reverse orientation is eligible to be considered in the upstream (downstream) path.

As an example, let us assume in Figure 2, *A*^+^ denotes the returned AMR gene path in graph by the sequence alignment tool (i.e., Bandage-BLASTN or GraphAligner). To extract the upstream, we can include node *B*^−^ because the orientation of *A* in edge *B*^−^ →*A*^+^ is the same as the orientation of *A* in the returned AMR gene path. However, as shown in Figure 2a, it seems that the sequence of no further nodes in the upstream path can be added as there is no edge from any other node to *B*^−^. If we take advantage of the idea of reversing edges, *B*^+^→ *C*^−^ can be converted to *C*^+^→ *B*^−^, and as shown in Figure 2b, the sequence of *C*^+^ can be added to the upstream path as well.

**Figure 2.**
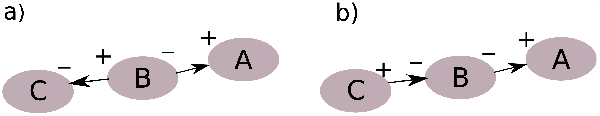
Utilizing reverse edges in the neighborhood extraction process. (a)No path between *A* and *C* can be found by using the edges as they are available in the assembly graph, **(b)** by reversing the edge between *B* and *C* and converting *B*^+^ → *C*^−^ to *C*^+^ → *B*^−^, we can make a path from *C* to *A*.

Each extracted sequence, representing a graph path, is added to the list of extracted sequences if there is no similar sequence to it available in the list. If a similar sequence exists but it is shorter, we replace it with the new sequence. We extract the path for each neighborhood sequence, including a list of nodes, parts of the sequence represented by each node, and each node’s coverage. Figure 3 illustrates an example of neighborhood extraction.

**Figure 3.**
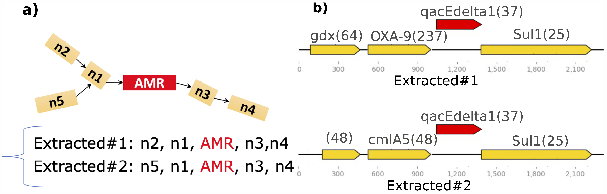
Neighborhood extraction. **(a)** The neighborhood of an AMR gene (*qacEdelta1*) is extracted as two separate paths with a maximum length of 1000 bp on each side of the AMR gene. **(b)** Two neighborhood sequences are annotated with the numbers in parentheses presenting gene coverage. Since the ratio of the reads matching *OXA-9* (i.e., 237) to the *qacEdelta1* gene (i.e., 37) exceeds a given threshold (i.e., 30), *OXA-9* and any gene in its upstream are removed resulting in removing the corresponding extracted upstream sequence from the candidate path list.

### Comparing annotations

After extracting neighborhood sequences for each AMR gene, we run Bakta or Prokka to identify gene information including name, start and end positions in the sequence, length, and locus tag. Annotated protein sequences are then processed using CARD’s Resistance Gene Identifier (RGI) to recognize any potential AMR gene; any strict/perfect annotations from RGI then supersede those identified by Bakta or Prokka. Only unique annotations are retained for different paths; by default, two annotations with the same number of genes are considered identical only if they share an identical annotation or have a pairwise BLASTN identify and coverage greater than 90%.

### Gene coverage and path filtering

In many cases, we expect the graph to contain false-positive chimeric paths (i.e., false pairings of upstream and downstream neighbourboods which do not exist within the underlying genomes) which require filtering. To reduce the number of such paths, we consider the relative abundance or *coverage* of each node, calculated as:

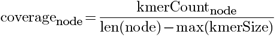

where *kmerCount*_*node*_ is the number of times k-mers of the node appear in the reads, *len*(*node*) is the length of the node in nucleotides, and *max*(*kmerSize*) is the maximum k-mer size used by the assembler.

The coverage for each annotated gene is a weighted average over the coverage of all nodes covering the gene sequence, with the weight of each node expressed as the length of the gene sequence contained in the node divided by the total sequence length. For example, for a gene, with length *L*, represented as path *N*_1_ →*N*_2_ →*N*_3_, where its *L*_1_, *L*_2_ and *L*_3_ nucleotides are extracted from nodes *N*_1_, *N*_2_ and *N*_3_, respectively, the coverage equals:

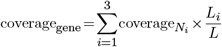

We use a coverage ratio threshold between the target AMR gene and other genes in each annotated extracted sequence to remove implausible paths. If the ratio of coverage between gene *G* and the target AMR gene exceeds this threshold, then gene *G* and any other gene(s) in its upstream or downstream path are removed. This process has been illustrated with an example in Figure 3b, where gene *OXA-9* and all its upstream genes are removed from the extracted sequence due to the considerable difference in coverage between *OXA-9* and the target AMR gene (i.e., *qacEdelta1*). All remaining unique annotated sequences are returned by Sarand and visualized.

### Validation of Sarand

We ran Sarand (v1.0.0) with GFApy v1.1.0 (39), Bandage v0.8.1 (31), Prokka v1.14.5 (34), RGI v5.1.1 (38), CARD v3.1.0 (30) and dna features viewer v3.0.3 (35). As presented in Table 1, we validated our software on three classes of datasets: (i) simple metagenomes constructed from either one or two strains from each of *Escherichia coli, Staphylococcus aureus*, and *Klebsiella pneumoniae* retrieved from RefSeq, designated as ‘1_1_1’ and ‘2_2_2’ respectively, with reads simulated using ART V2.5.8 (40) (HiSeq 2500 with read length=150bp, insert size=500bp, fold coverage=20); (ii) “medium” and “high”-complexity datasets from the Critical Assessment of Metagenome Interpretation (CAMI) study (41); and (iii) three published metagenome samples (28) derived from urban sewage sequenced by Illumina HiSeq.

**Table 1.**
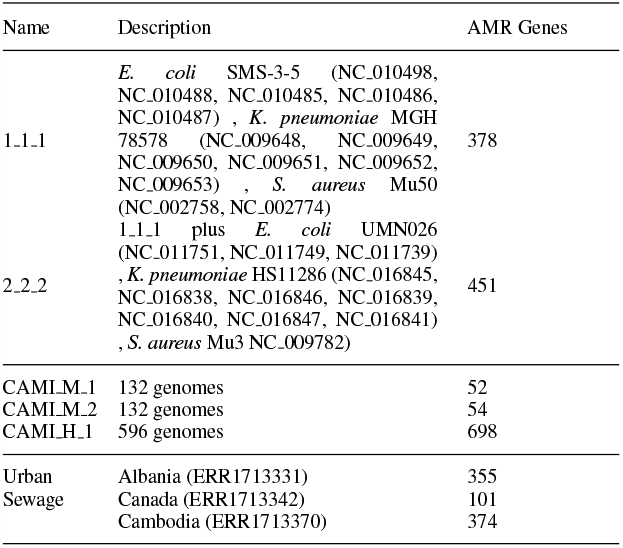
Datasets simulated from RefSeq (1_1_1, 2_2_2), the CAMI Challenge (42), and real metagenomic samples selected from the Global Urban Sewage AMR Monitoring Project (28).

We examined the sensitivity and precision of Sarand on the simulated datasets (i) and (ii). To evaluate the validity of extracted sequences, we compared them with the ground truth available in simulated datasets. For datasets 1_1_1 and 2_2_2, we used BLASTN v2.9.0 to query the AMR genes on the reference genomes (the default threshold for coverage and identity was set to 95%) and then extracted their neighborhood sequences to be compared with Sarand’s extracted neighborhoods. For the CAMI dataset, extracted neighborhoods passing the Sarand thresholds were similarly compared against the gold-standard FASTA files available from the CAMI website. If an upstream (or downstream) sequence from the assembly graph matched true upstream (or downstream) sequences from the reference genomes, it was considered as a true positive; extracted sequences with no match in the ground truth were counted as false positives.

We compared the accuracy of Sarand against contigs assembled by metaSPAdes v3.14.1 (36) as well as two recently developed tools representing two different paradigms, MetaCherchant (25) and Spacegraphcats (26). For MetaCherchant v0.1.0, we used the default settings (*k*=31, maxkmers=100000, bothdirs=false, chunklength=10) with coverage=1 to include as many k-mers as possible in the results as recommended by the developers (*pers. comm*.). We applied a modified version of our method to extract all available neighborhood sequences from the MetaCherchant graph. In most cases, it did not return any path because either the number of nodes representing the AMR sequences in the MetaCherchant graph was greater than 50 (i.e., the maximum path length used in Bandage+BLAST queries), or there was no continuous path of nodes, representing AMR sequence, available in the graph. To address this limitation, we found the order of the nodes representing the AMR gene by aligning their sequences over the AMR sequence and ran our recursive functions to extract the upstream and downstream sequences.

We ran Spacegraphcats v2.0.12 for datasets 1_1_1 and 2_2_2 with radius=10 and *k*=31. However, for CAMI datasets and radius > 1, we exceeded our 1.4TB available memory. Therefore, for CAMI datasets, we were only able to run Spacegraphcats with radius=1. For each dataset, Spacegraphcats constructs a de Bruijn graph by running BCALM. After processing this graph, for each query (AMR sequence), it returns a list of nodes that contain the query neighborhood. We first extracted a subgraph of the original BCALM graph containing the output nodes as well as the nodes which are one hop away from them. Then, a modified version of our method was applied to extract the neighborhood sequences of the target AMR gene from the subgraph, filter and annotate them. However, in almost all cases, the length of the extracted neighborhood sequence was not long enough to be annotated as any gene(s).

All experiments were run on a computer workstation running Ubuntu 20.04 with 3.5 GHZ Intel Xeon Quad-Core Processor and 128 GB RAM, except for CAMI experiments with Spacegraphcats that required a significant amount of memory and were run on Ubuntu 20.04 server with 88 Xeon Processor cores and 1.4 TB memory.

## RESULTS

### Performance on simulated datasets

#### Identification of AMR genes

To establish a performance baseline and ground truth, we compared the sensitivity of BLASTN for putative AMR gene detection in the underlying reference genomes, metaSPAdes assembly graphs (using Bandage+BLASTN), and metaSPAdes contigs across the simulated datasets (Table 2). Although nearly all AMR genes identified in the underlying reference genomes were also detected in the assembly graphs and contigs, there was a general trend of decreasing numbers of detectable AMR genes (reference genome > assembly graph > contigs) across all datasets. This supports our hypothesis that some AMR sequences are lost in the process of extracting contigs from the assembly graph.

**Table 2.**
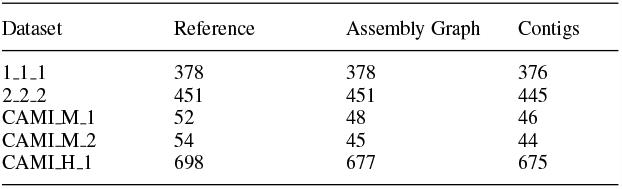
AMR genes identified by BLASTN in each dataset.

| Dataset | Reference | Assembly Graph | Contigs |
| --- | --- | --- | --- |
| 1_1_1 | 378 | 378 | 376 |
| 2_2_2 | 451 | 451 | 445 |
| CAMI_M_1 | 52 | 48 | 46 |
| CAMI_M_2 | 54 | 45 | 44 |
| CAMI_H_1 | 698 | 677 | 675 |

#### Extraction of AMR neighborhoods

Using the underlying reference genomes as the ground-truth, we then evaluated the performance of Sarand, MetaCherchant, and metaSPAdes contigs in correctly extracting 1000bp upstream and downstream AMR gene neighborhoods. We did not include the results from Spacegraphcats because, as explained before, the neighbourhood sequences extracted by Spacegraphcats, for most datasets, were not long enough to represent any neighbourhood gene. We selected a relative gene coverage threshold of 30 for Sarand based on calibration against samples 1_1_1 and CAMI M 1 (Supplementary Figures 1 and 2). Figures 4a and 4b show the precision and sensitivity of these 3 different methods. Sarand drastically outperformed MetaCherchant and the contig-based approach on the simplest datasets, with near-perfect sensitivity and 76% specificity on the 1_1_1 dataset, with contig precision the only other score in excess of 50%. Adding a second genome of each of the three species led to a drastic reduction in accuracy: while the precision of Sarand remained relatively high at 87%, the sensitivity dropped to 34%. The performance of both other methods uniformly decreased as well. When applied to the CAMI datasets, Sarand remained the top-performing method in terms of precision and sensitivity, albeit with much smaller margins relative to the contig-based measures. Consistent with the results on the simple simulated datasets, the performance of all methods tended to decrease with increasing complexity (i.e., from CAMI M 1 to CAMI M 2 to CAMI H 1), with the notable exception of MetaCherchant which performed best on CAMI M 2 in terms of both precision and sensitivity.

**Figure 4.**
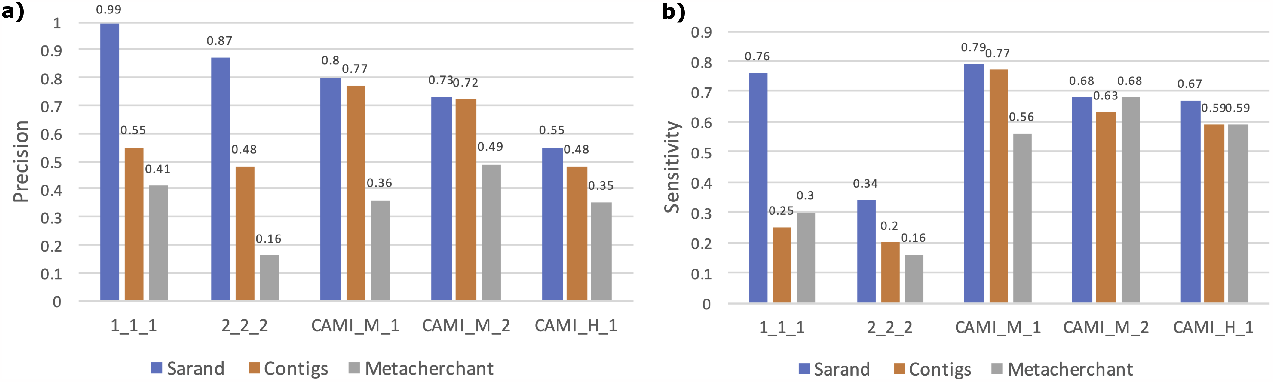
Performance comparison across simulated datasets. The gene-coverage threshold for filtering in Sarand was set to 30. **(a) Average precision**. Precision measures the fraction of extracted upstream/downstream annotations that are present in the reference genomes. **(b) Average sensitivity**. Sensitivity measures the fraction of upstream/downstream annotations from the reference that were identified by a method (i.e., Sarand, MetaCherchant, or contigs).

*sul1* in the CAMI H 1 sample is an example case where Sarand outperforms contigs and MetaCherchant. As shown in Figure 5, Sarand successfully reconstructed two neighbourhoods, each of which was consistent with one of the two reference sequences. The contig-based approach reconstructed only one of the two references, which was also identified by MetaCherchant. However, MetaCherchant also made two incorrect predictions, each of which contained false-positive genes that were absent from the reference. Conversely, Figure 6 illustrates one of the cases where Sarand, contigs, and MetaCherchant all fail to identify the correct neighborhood. For *APH(3’)-Ia* in CAMI H 1, Sarand only identifies an annotated downstream sequence which is not present in the reference genome. The correct upstream gene was correctly identified by Sarand but then removed by the coverage filter.

**Figure 5.**
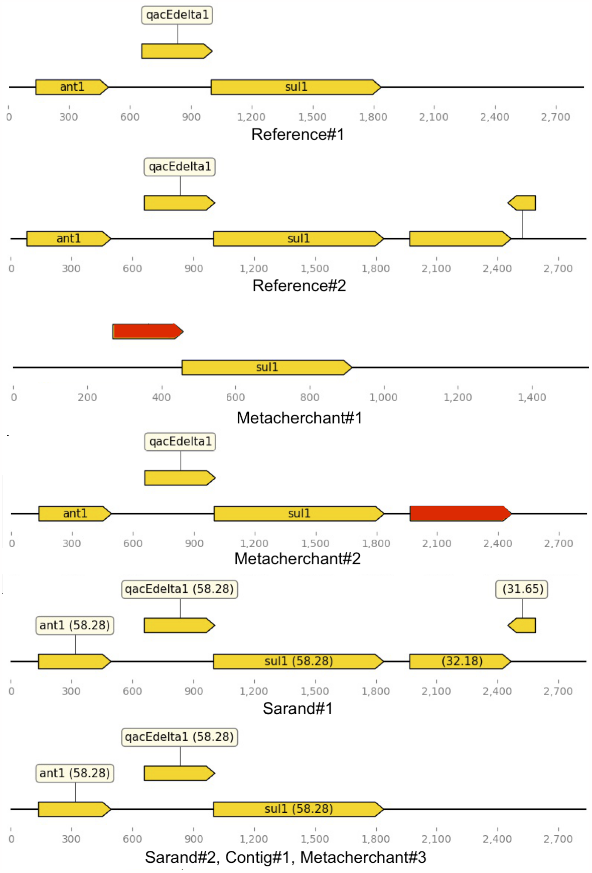
The annotation of neighborhood sequences extracted for *sul1*, from CAMI H 1, by Sarand, MetaCherchant and contigs. Please note that Sarand#2, Contig#1 and MetaCherchant#3 only include the upstream sequence, and false-positive genes are highlighted in red.

**Figure 6.**
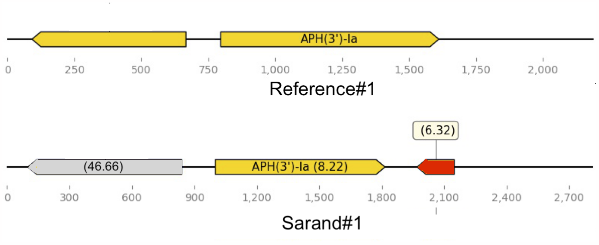
The annotation of neighborhood sequences extracted for *APH(3’)-Ia*, from Sarand and a reference long contig in the CAMI H 1 dataset. The gene highlighted in red is a false-positive case while the gene highlighted in grey was initially identified by Sarand but removed after applying Sarand’s differential gene coverage statistic.

We then compared the number of AMR genes whose upstream and downstream neighborhoods were correctly extracted across each method and dataset. This was summarised across 2 different sensitivity thresholds: number of AMR genes with all their upstream and downstream neighborhoods detected successfully, and number of AMR genes with at least half of their upstream and downstream neighborhoods detected successfully (Supplementary Figures 3-12).

As shown in these figures, for samples 1_1_1 and 2_2_2, Sarand was able to fully reconstruct the neighborhood of more AMR genes than contigs and MetaCherchant. Also, it identified at least 50% of the true neighborhoods for 86% of AMR genes in 1_1_1 and 37% of AMR genes in 2_2_2 that were detected by neither contigs nor MetaCherchant. In the CAMI datasets, MetaCherchant was able to reconstruct the most full neighborhoods (2%, 11% and 23% of AMR genes for CAMI M 1, CAMI M 2 and CAMI H 1, respectively). However, as shown in Figure 4b, when comparing across all simulated samples, Sarand identifies neighborhoods with higher sensitivity than MetaCherchant and contigs.

### Detection of AMR gene neighbourhoods in sewage samples

We applied Sarand to three published metagenome samples from urban sewage sequenced using the Illumina HiSeq (28). We chose three published samples from disparate geographic locations with different numbers and types of predicted AMR genes: sample ERR1713331 from Albania (355 AMR genes), sample ERR1713342 from Canada (101 AMR genes), and sample ERR1713370 from Cambodia (374 AMR genes).

After assembly in metaSPAdes, we ran Sarand for the constructed assembly graph of each sample and extracted the list of filtered neighborhood sequences of each AMR gene, based on a gene-coverage ratio threshold of 30, as explained in Section “Materials and Methods”.

#### Presence of extracted sequences in reference databases

Although we anticipate that previously undiscovered AMR gene neighbourhoods will be present in complex, global metagenome samples, we can still assess the performance of Sarand in recovering neighborhoods that are already present in reference databases. To assess this, we searched the neighborhoods extracted by Sarand against the NCBI ‘nt’ database (August 2021) using BLASTN with a percent-identity threshold of 90%. We classified hits as “full match” if the entire predicted neighborhood of a given AMR gene was present in the database, “upstream+AMR” or “AMR+downstream” if only the upstream or downstream neighborhoods predicted by Sarand, along with the AMR sequence, were found in the database, or “no hit” if neither the upstream nor downstream predicted neighborhoods were present.

The summary of the best BLAST hit results across all extracted neighborhood sequences for every samples is shown in Figure 7. In general, for sample *ERR1713331*, about 56% of the extracted neighborhood sequences have at least partial matches, with more than half of “no match” sequences belonging to the OXA beta-lactamase family. In samples *ERR1713342* and *ERR1713370* about three quarters of the extracted sequences had at least partial (i.e., upstream+AMR or AMR+downstream) matches, although full matches were fewer in number than either upstream or downstream matches.

**Figure 7.**
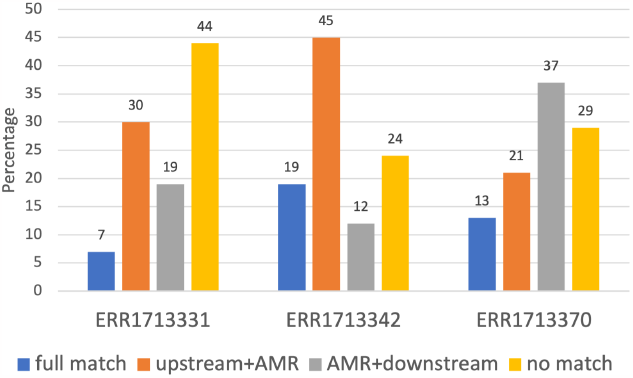
Summary results of BLAST hits for all neighborhood sequences extracted from sewage samples. BLASTN was used to search the neighborhoods extracted by Sarand against NCBI ‘nt’ database and the results were classified as “full match” (the entire sequence is present in the database), “upstream+AMR” (only upstream sequence is present), “downstrean+AMR” (only downstream sequence is present) and “no hit” (neither upstream nor downstream is present).

## DISCUSSION

This study, in line with other recent research (25, 26, 43), showed that making gene-neighbourhood predictions directly from the assembly graph can be more informative than exclusively using contigs. Comparing AMR neighborhoods across all 5 different simulated datasets used in this study, Sarand performs on average 47% and 46% better than assembled metagenomic contigs in terms of precision and sensitivity, respectively. Similarly, we also showed through the 134% precision and 48% sensitivity improvement over MetaCherchant that methods making use of the full metagenomic assembly graph are better able to capture gene neighbourhoods than local assemblies. Unfortunately, we were not able to fully evaluate Spacegraphcats because for larger datasets, such as CAMI samples, it was too memory-intensive to run with comparable neighbourhood sizes (i.e., with radius> 1). It may be possible to improve the performance of the dominating sets approach for this specific problem. This could involve more aggressive denoising of the assembly graph and additional heuristics more optimised to individual gene scale neighbourhoods rather than the genome scale queries for which spacegraphcats was developed.

Simulation verisimilitude is a common limitation in any benchmarking analysis using simulated data. However, without the reliable ground-truth provided by simulation, it is often challenging to fully evaluate a tool’s performance. We attempted to address this through our analysis of wastewater metagenomic sequencing data (with 56-76% of Sarand neighbourhoods identifiable in NCBI deposited genomes) but further work could include sequenced mock microbial communities (e.g., ZymoBIOMICS Microbial Community Standards). We also only evaluated one homology-search paradigm (i.e., Bandage’s BLAST implementation) and one input metagenomic graph (i.e., MetaSPAdes) despite Sarand supporting GraphAligner and MegaHit or BCALM graphs. A formal evaluation and comparison of the many emerging graph homology search methods (e.g., Wheeler graph-based methods (44), hidden Markov models (45), and k-mer methods (46)) would be a valuable resource to tool developers. Finally, despite the increased availability of high depth long-read metagenomic data, we did not evaluate how effectively Sarand adapts to the long-read metagenomic assembly graphs.

## CONCLUSION

Sarand combines the sensitivity of read-based approaches with the genomic context provided as assemblies to profile AMR genes and their associated gene neighboods from metagenomic samples. Across several simulated datasets, we were able to show that Sarand had a higher precision and sensitivity than alternative methods (e.g., MetaCherchant, metaSPAdes assembly, and spacegraphcats). Future work will seek to extend the coverage threshold approach to further reduce chimeric neighborhoods (false joining of upstream and downstream neighborhoods), incorporate alternative homology search algorithms, and support long-read datasets.

## ACKNOWLEDGEMENTS

SK was supported by Saint Mary’s University and CIHR Project Grant funding held by RGB. FM was supported by a Donald Hill Family Fellowship in Computer Science and an NSERC Discovery Grant. Computer resources were supplied by the McMaster University Service Lab and Repository computing cluster, courtesy of Andrew McArthur (Department of Biochemistry & Biomedical Sciences), funded in part by grants from the Canada Foundation for Innovation and a donation of hardware by Cisco Systems Canada, Inc.

