## Supplemental File for "Sarand: Exploring Antimicrobial Resistance Gene Neighborhoods in Complex Metagenomic Assembly Graphs"

### A - CHOOSING GENE COVERAGE THRESHOLD

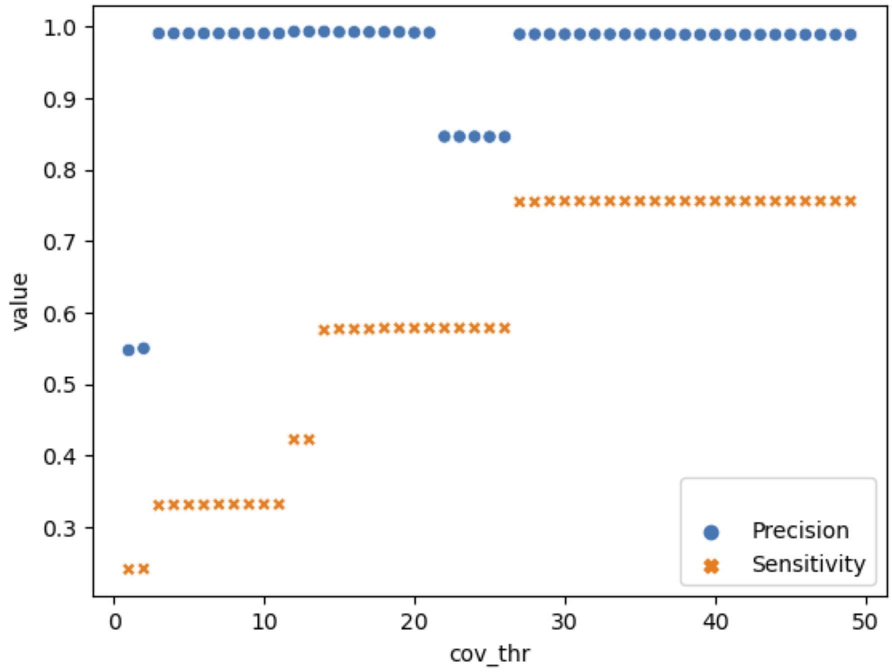


**Figure 1.** Precision and sensitivity of sample 1_1_1 for extracted neighborhood sequences across different relative gene-coverage threshold values.

© The Author(s)
This is an Open Access article distributed under the terms of the Creative Commons Attribution Non-Commercial License (http://creativecommons.org/licenses/by-nc/2.0/uk/) which permits unrestricted non-commercial use, distribution, and reproduction in any medium, provided the original work is properly cited.

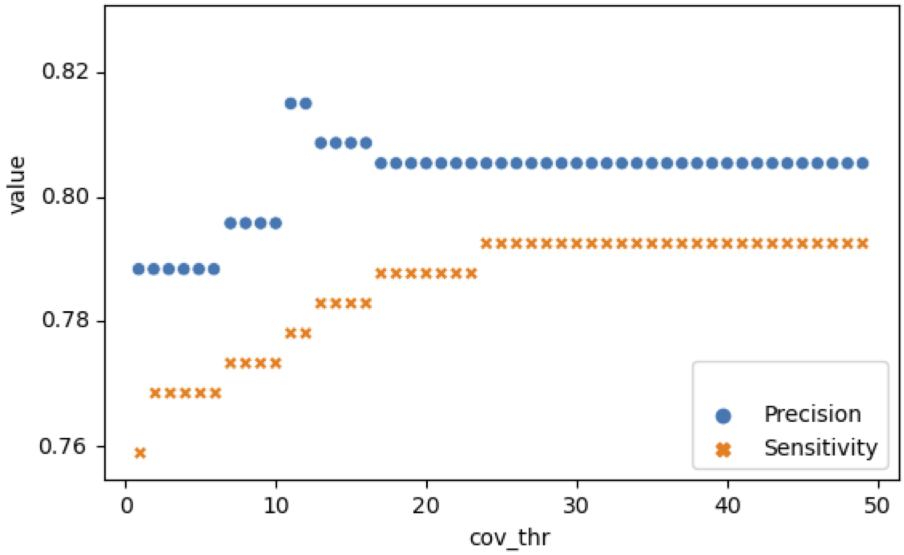


**Figure 2.** Precision and sensitivity of sample CAMI_M_1 for extracted neighborhood sequences across different relative gene-coverage threshold values.

### B - COMPARISON OF CORRECTLY DETECTED AMR NEIGHBORHOODS FOR DIFFERENT DATASETS

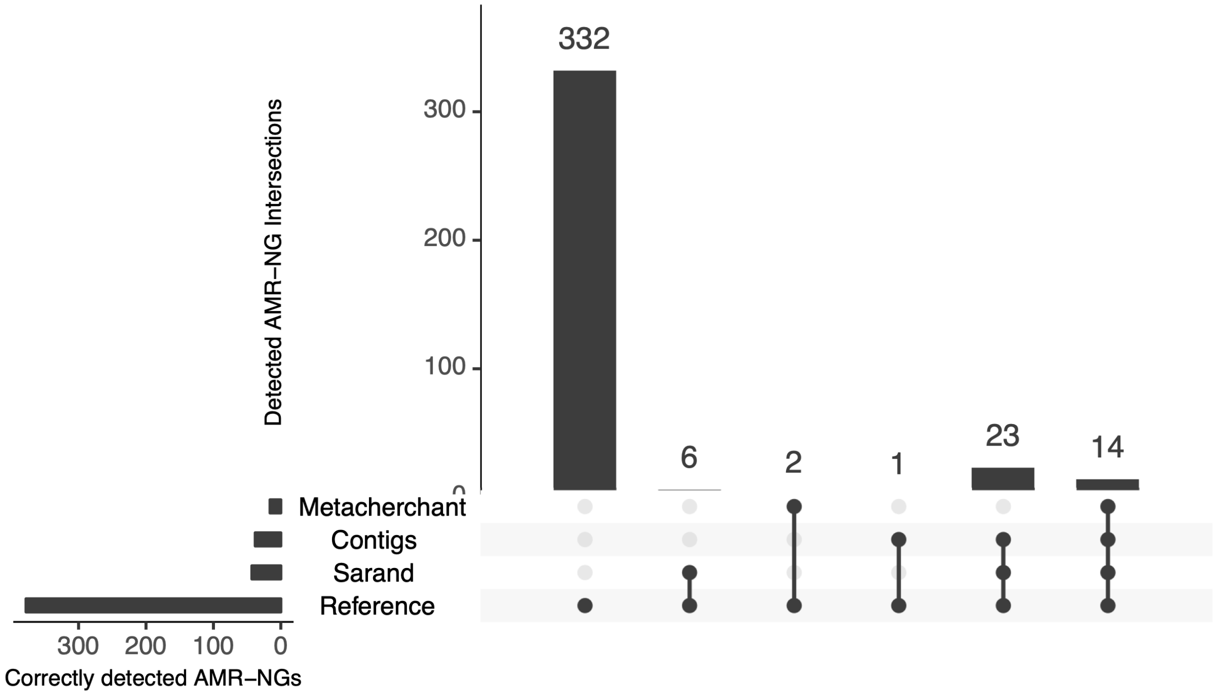


**Figure 3.** Comparison of correctly detected AMR neighborhoods with sensitivity = 1 for 1_1_1 dataset.

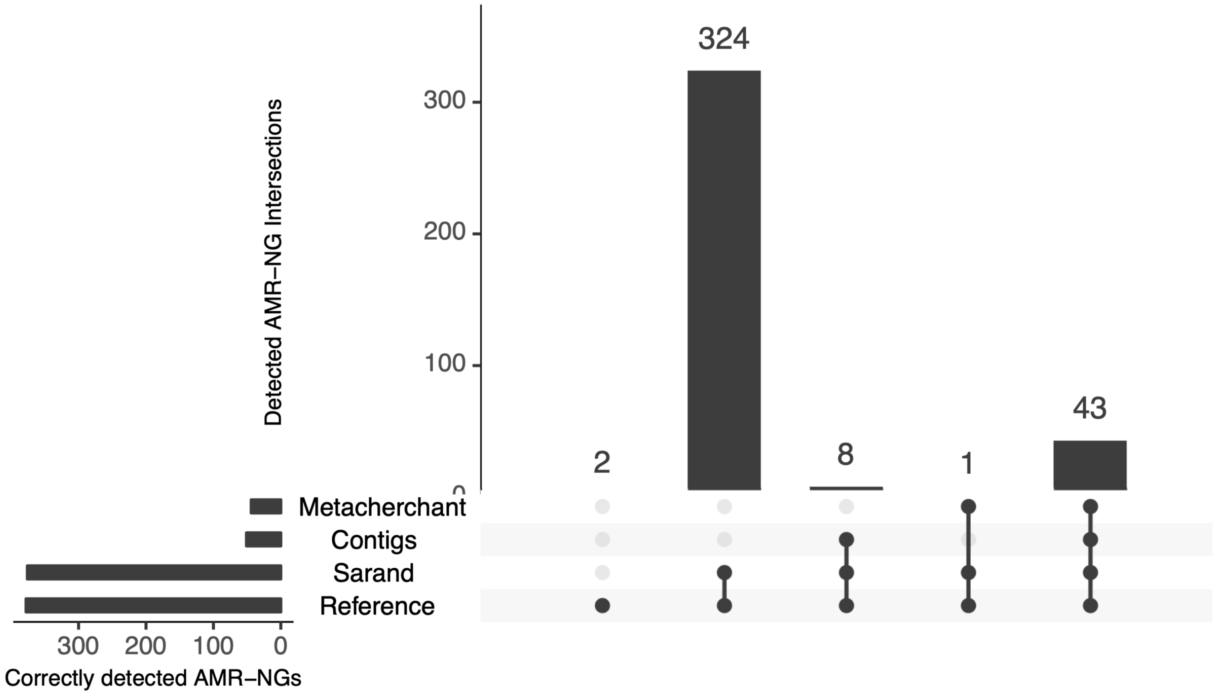


**Figure 4.** Comparison of correctly detected AMR neighborhoods with sensitivity = 0.5 for 1_1_1 dataset.

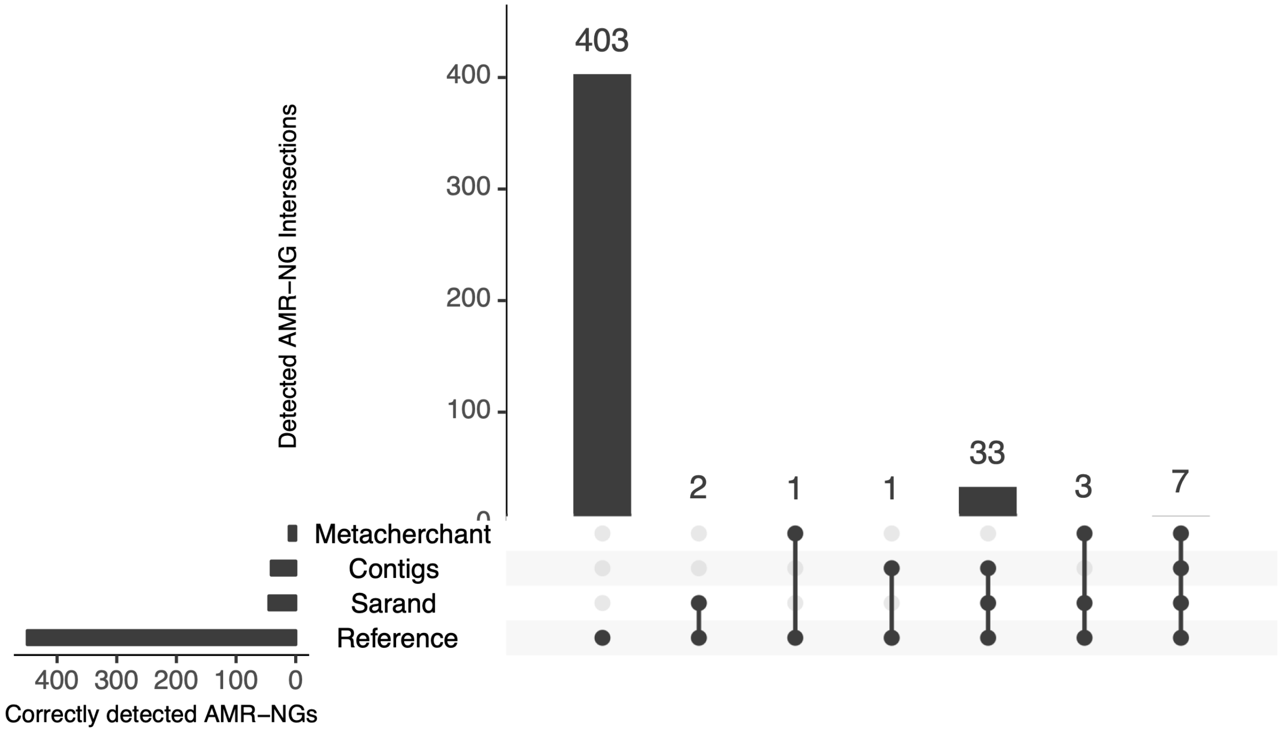


**Figure 5.** Comparison of correctly detected AMR neighborhoods with sensitivity = 1 for 2_2_2 dataset.

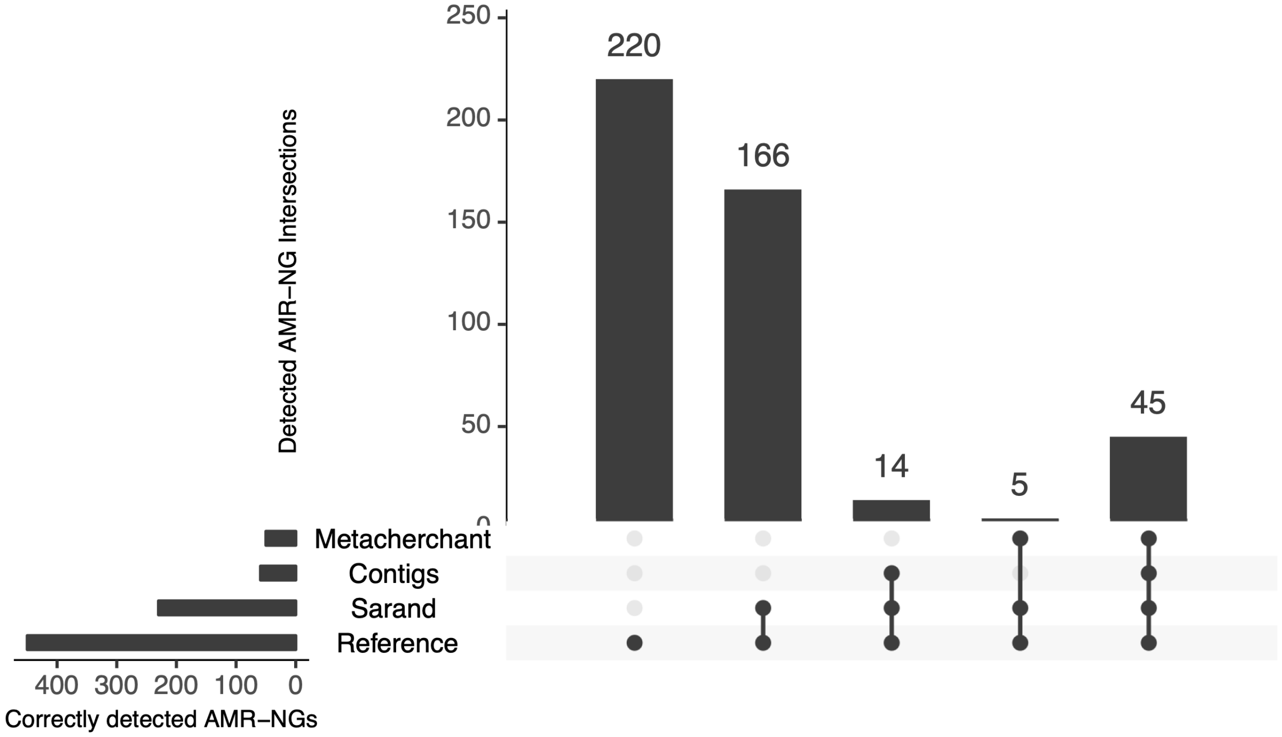


**Figure 6.** Comparison of correctly detected AMR neighborhoods with sensitivity = 0.5 for 2_2_2 dataset.

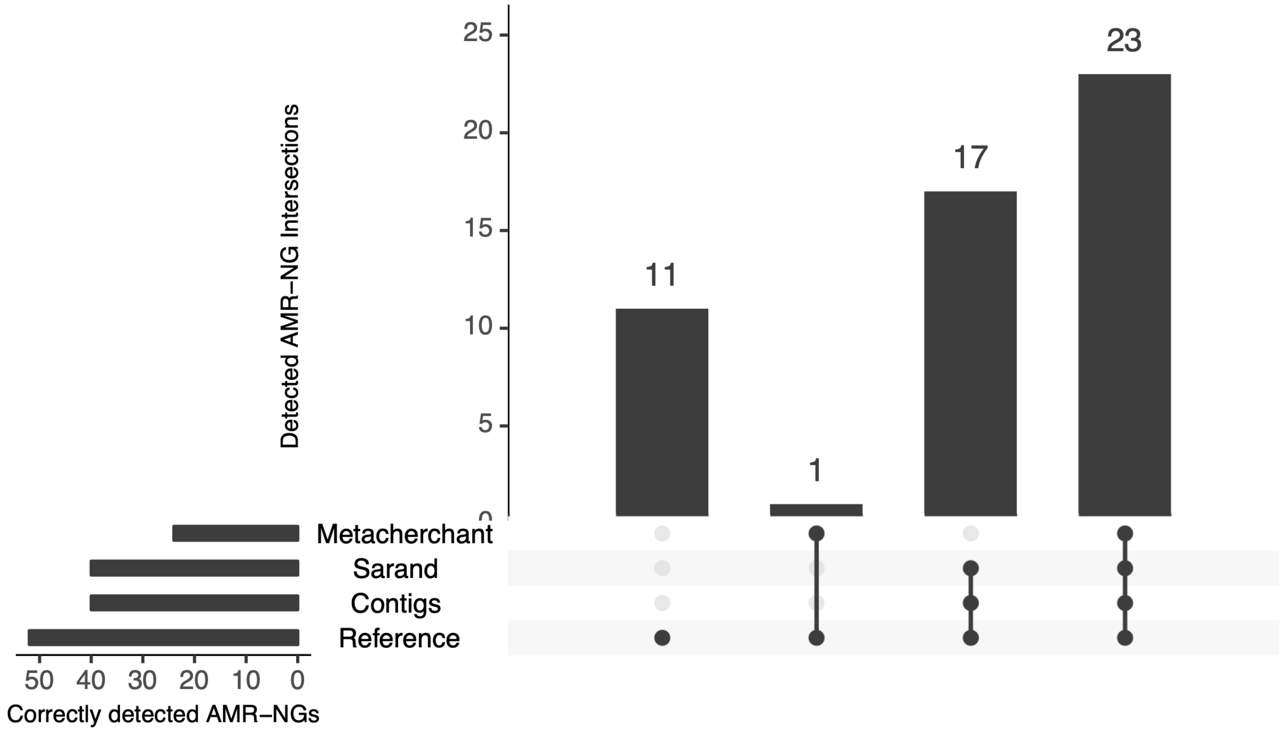


**Figure 7.** Comparison of correctly detected AMR neighborhoods with sensitivity = 1 for CAMI_M_1 dataset.

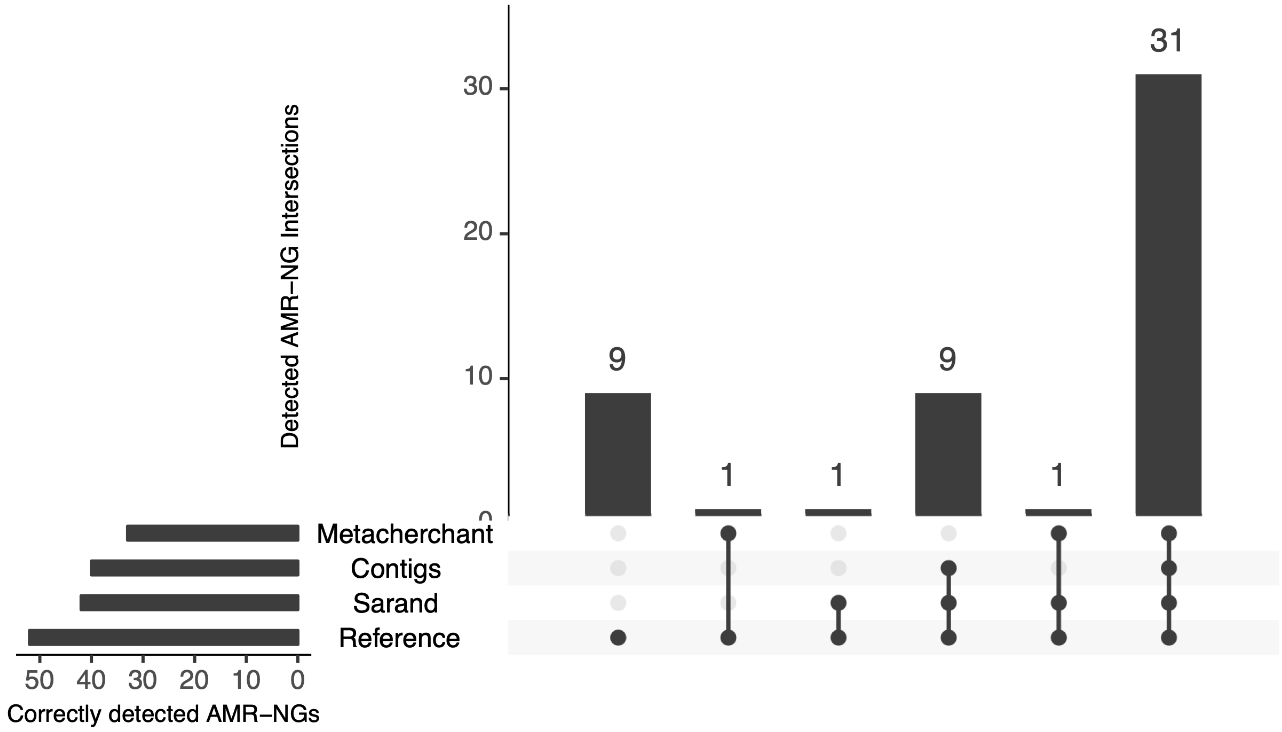


**Figure 8.** Comparison of correctly detected AMR neighborhoods with sensitivity = 0.5 for CAMI_M_1 dataset.

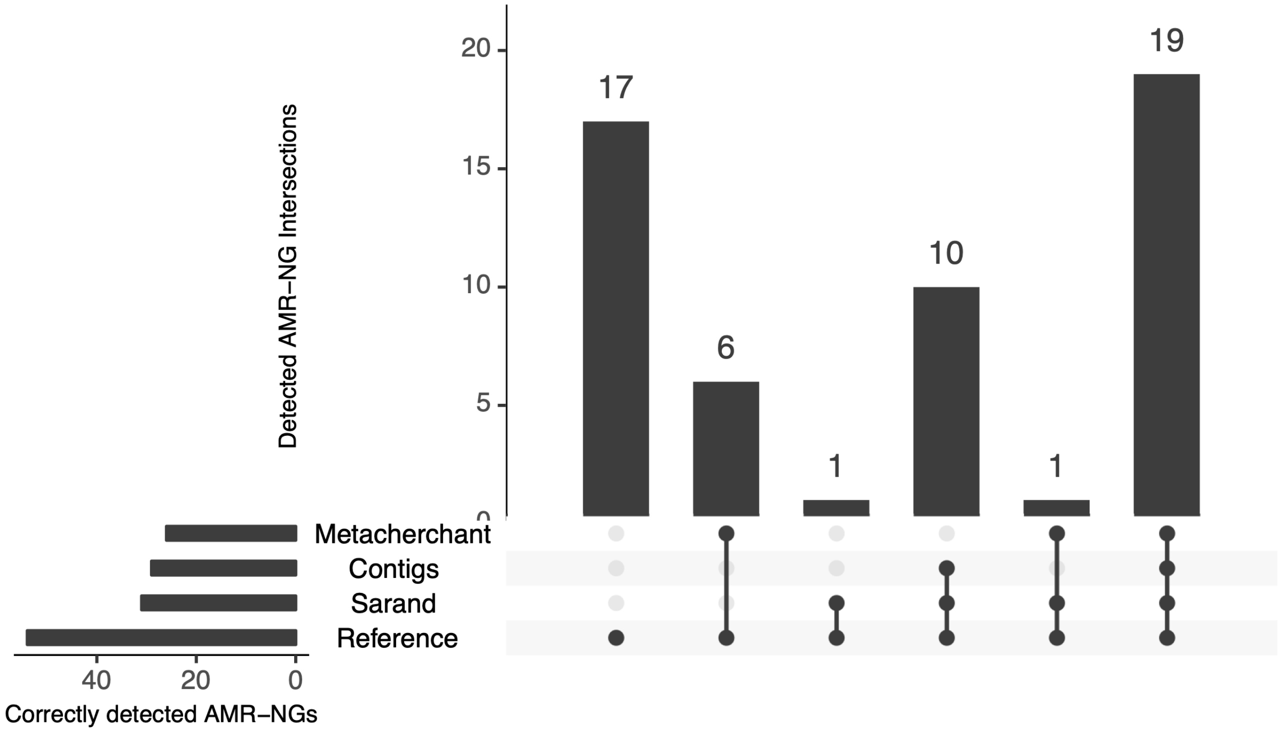


**Figure 9.** Comparison of correctly detected AMR neighborhoods with sensitivity = 1 for CAMI_M_2 dataset.

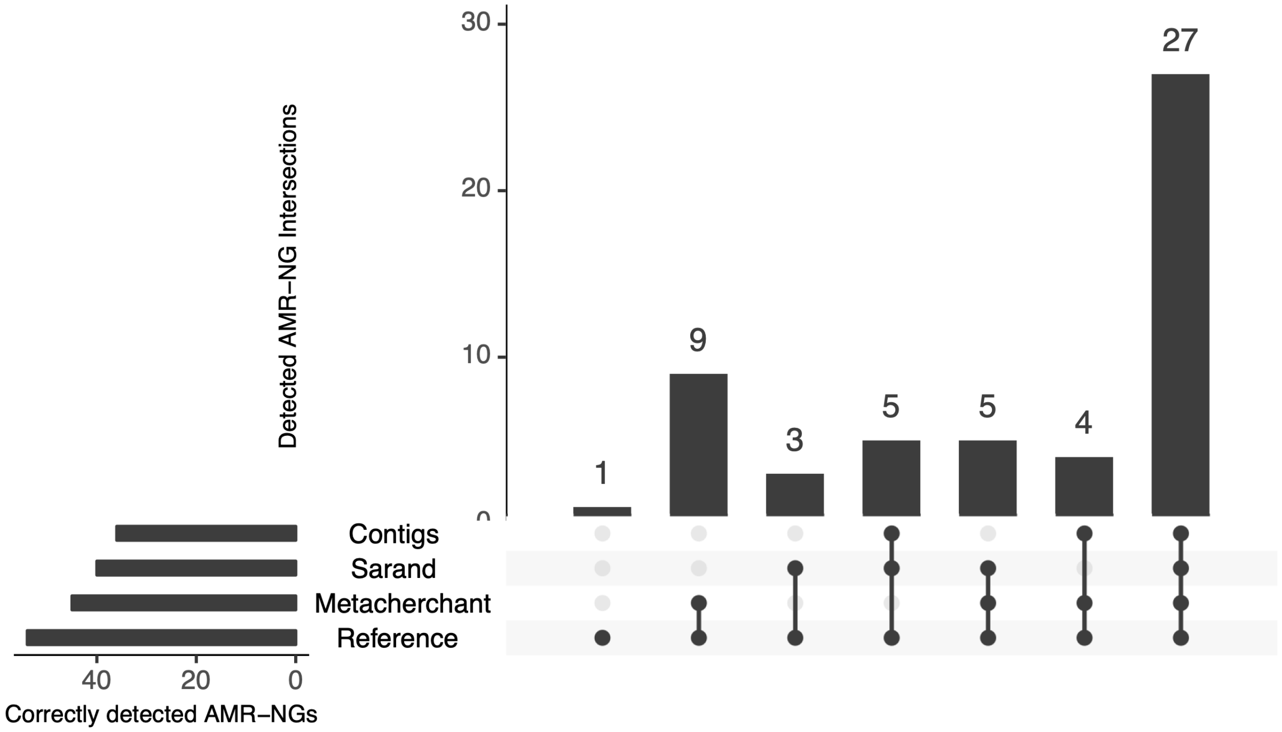


**Figure 10.** Comparison of correctly detected AMR neighborhoods with sensitivity = 0.5 for CAMI_M_2 dataset.

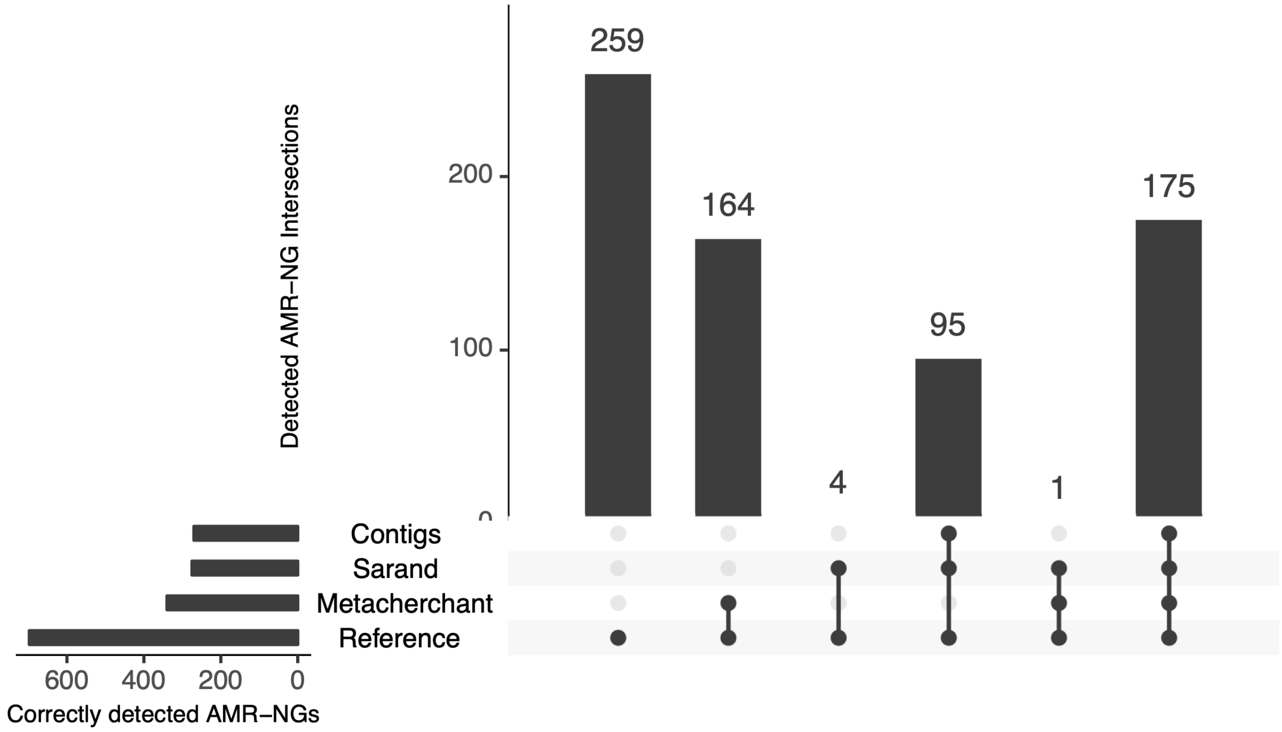


**Figure 11.** Comparison of correctly detected AMR neighborhoods with sensitivity = 1 for CAMI_H_1 dataset.

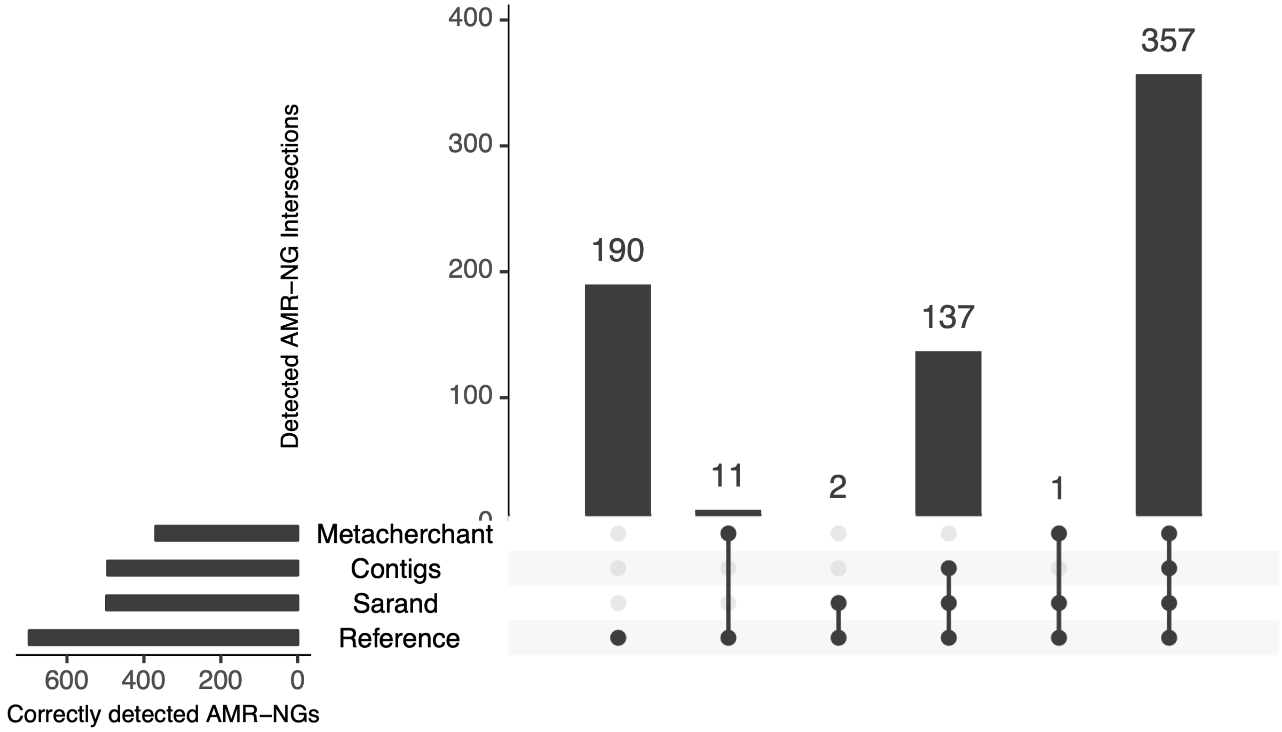


**Figure 12.** Comparison of correctly detected AMR neighborhoods with sensitivity = 0.5 for CAMI_H_1 dataset.

### C - SARAND PERFORMANCE EVALUATION - EXCEPTIONS

**Table 1.** All cases that contig outperforms Sarand. P and S refer to precision and sensitivity, respectively.

| Dataset | AMR | Contig (P, S) | Sarand (P, S) |
|---|---|---|---|
| 1_1_1 | APH(6)-Id | (1, 0.33) | (0.75, 1) |
| 1_1_1 | OXA-9 | (1, 0.5) | (0.67, 1) |
| 1_1_1 | Staphylococcus_aureus_FosB | (1, 1) | (0.5, 0.5) |
| 1_1_1 | sul1 | (1, 1) | (0.5, 1) |
| 2_2_2 | AAC(6’)-Ib, AAC(6’)-Ib-cr, AAC(6’)-Ib-Hangzhou, AAC(6’)-Ib-Suzhou, AAC(6’)-Ib’, AAC(6’)-Ib10, AAC(6’)-Ib11, AAC(6’)-Ib3, AAC(6’)-Ib4, AAC(6’)-Ib9 | (1, 0.5) | (0.5, 0.5) |
| 2_2_2 | APH(6)-Id | (1, 0.25) | (0.33, 0.5) |
| 2_2_2 | OXA-9 | (1, 0.5) | (0.5, 0.5) |
| 2_2_2 | Staphylococcus_aureus_FosB | (1, 1) | (0.5, 0.5) |
| CAMI_M_2 | aadA, aadA21, aadA22, ANT(3”)-IIa | (1, 0.5) | (0, 0) |
| CAMI_H_1 | aadK | (1, 1) | (0, 1) |
| CAMI_H_1 | catB9 | (1, 1) | (0.08, 1) |
| CAMI_H_1 | cprS | (1, 1) | (0.67, 1) |
| CAMI_H_1 | dfrA6, dfrA31 | (1, 0.5) | (0.5, 0.5) |
| CAMI_H_1 | sul3 | (1, 0.5) | (0.5, 0.5) |
| CAMI_H_1 | tetO | (1, 1) | (0.5, 1) |

**Table 2.** All cases that Metacherchant outperforms Sarand. P and S refer to precision and sensitivity, respectively.

| Dataset | AMR | Metacherchant (P, S) | Sarand (P, S) |
|---|---|---|---|
| 1_1_1 | AAC(6')-Ie-APH(2")-Ia | (0.67, 1) | (1, 0.5) |
| 1_1_1 | catII_from_Escherichia_coli_K-12 | (0.4, 1) | (1, 0.5) |
| 2_2_2 | catII_from_Escherichia_coli_K-12 | (0.67, 1) | (1, 0.5) |
| 2_2_2 | aadA12 | (0.5,0.5) | (0.25,0.5) |
| 2_2_2 | sul1 | (0.2,0.33) | (0.14,0.67) |
| CAMI_M_1 | OXA-724 | (0.25, 1) | (0, 0) |
| CAMI_M_1 | tet(W/N/W) | (0.5, 0.25) | (0, 0) |
| CAMI_M_2 | Escherichia_coli_ampC1_betalactamase | (0.5, 1) | (0.5, 0.5) |
| CAMI_M_2 | AAC(6')-Ii, eptA, kdpE | (1, 1) | (0, 0) |
| CAMI_M_2 | ErmG | (0.25, 1) | (0, 0) |
| CAMI_M_2 | tet32 | (0, 1) | (0, 0) |
| CAMI_M_2 | aadA, aadA15, aadA21, aadA22, aadA23, aadA24, ANT(3")-IIa | (1, 0.5) | (0, 0) |
| CAMI_M_2 | bacA | (0.33, 0.5) | (0, 0) |
| CAMI_H_1 | Escherichia_coli_ampC1_betalactamase | (0.67, 1) | (0.5, 0.5) |
| CAMI_H_1 | FosB | (0.25, 1) | (0, 0) |
| CAMI_H_1 | OXA-429, OXA-430, OXA-431, OXA-432, OXA-433, OXA-441, OXA-442, OXA-480, OXA-51, OXA-64, OXA-65, OXA-66, OXA-67, OXA-68, OXA-69, OXA-70, OXA-71, OXA-75, OXA-76, OXA-77, OXA-174, OXA-378, OXA-426, OXA-78, OXA-175, OXA-176, OXA-177, OXA-178, OXA-179, OXA-180, OXA-194, OXA-195, OXA-196, OXA-197, OXA-200, OXA-201, OXA-202, OXA-203, OXA-206, OXA-208, OXA-100, OXA-106, OXA-107, OXA-108, OXA-109, OXA-110, OXA-111, OXA-112, OXA-113, OXA-115, OXA-116, OXA-117, OXA-120, OXA-121, OXA-122, OXA-123, OXA-124, OXA-125, OXA-126, OXA-127, OXA-128, OXA-130, OXA-131, OXA-132, OXA-312, OXA-313, OXA-314, OXA-315, OXA-316, OXA-317, OXA-336, OXA-337, OXA-338, OXA-339, OXA-340, OXA-341, OXA-342, OXA-343, OXA-344, OXA-345, OXA-346, OXA-365, OXA-371, OXA-374, OXA-375, OXA-376, OXA-377, OXA-138, OXA-144, OXA-148, OXA-149, OXA-150, OXA-172, OXA-173, OXA-79, OXA-80, OXA-82, OXA-83, OXA-84, OXA-86, OXA-87, OXA-88, OXA-89, OXA-90, OXA-91, OXA-92, OXA-93, OXA-94, OXA-95, OXA-98, OXA-99, OXA-216, OXA-217, OXA-219, OXA-223, OXA-234, OXA-241, OXA-242, OXA-248, OXA-249, OXA-250, OXA-254, OXA-259, OXA-260, OXA-261, OXA-262, OXA-263, OXA-379, OXA-380, OXA-381, OXA-382, OXA-383, OXA-384, OXA-385, OXA-386, OXA-387, OXA-388, OXA-389, OXA-390, OXA-391, OXA-400, OXA-401, OXA-402, OXA-403, OXA-404, OXA-406, OXA-407, OXA-408, OXA-409, OXA-411, OXA-412, OXA-413, OXA-414, OXA-424, OXA-425 | (0.75, 1) | (1, 0.67) |
| CAMI_H_1 | eptB | (0, 1) | (0, 0) |
| CAMI_H_1 | mphA, tmrB, ugd, vgaA, vgaALC | (1, 1) | (0, 0) |
| CAMI_H_1 | mphK | (0.5, 1) | (0, 0) |
| CAMI_H_1 | Bacillus_subtilis_mprF | (0.33, 0.5) | (0, 0) |
| CAMI_H_1 | BLA1 | (0.2, 0.5) | (0, 0) |
| CAMI_H_1 | clbA | (1, 0.5) | (0, 0) |
| CAMI_H_1 | aadK | (1, 1) | (0, 1) |
| CAMI_H_1 | SHV-1, SHV-101, SHV-102, SHV-103, SHV-104, SHV-105, SHV-106, SHV-107, SHV-108, SHV-109, SHV-11, SHV-110, SHV-111, SHV-119, SHV-12, SHV-120, SHV-121, SHV-122, SHV-123, SHV-124, SHV-125, SHV-126, SHV-127, SHV-128, SHV-129, SHV-13, SHV-133, SHV-134, SHV-135, SHV-137, SHV-14, SHV-140, SHV-141, SHV-142, SHV-143, SHV-144, SHV-145, SHV-147, SHV-148, SHV-149, SHV-15, SHV-150, SHV-151, SHV-152, SHV-153, SHV-154, SHV-155, SHV-156, SHV-157, SHV-158, SHV-159, SHV-16, SHV-160, SHV-161, SHV-162, SHV-163, SHV-164, SHV-165, SHV-167, SHV-168, SHV-172, SHV-173, SHV-178, SHV-179, SHV-18, SHV-180, SHV-182, SHV-183, SHV-185, SHV-186, SHV-187, SHV-188, SHV-189, SHV-19, SHV-2, SHV-20, SHV-21, SHV-22, SHV-23, SHV-24, SHV-25, SHV-26, SHV-27, SHV-28, SHV-29, SHV-2A, SHV-3, SHV-30, SHV-31, SHV-32, SHV-33, SHV-34, SHV-35, SHV-36, SHV-37, SHV-38, SHV-4, SHV-40, SHV-41, SHV-42, SHV-43, SHV-44, SHV-45, SHV-46, SHV-48, SHV-49, SHV-5, SHV-50, SHV-51, SHV-52, SHV-53, SHV-55, SHV-56, SHV-57, SHV-59, SHV-6, SHV-60, SHV-61, SHV-62, SHV-63, SHV-64, SHV-65, SHV-66, SHV-67, SHV-69, SHV-7, SHV-70, SHV-71, SHV-72, SHV-73, SHV-74, SHV-75, SHV-76, SHV-77, SHV-78, SHV-79, SHV-8, SHV-80, SHV-81, SHV-82, SHV-83, SHV-84, SHV-85, SHV-86, SHV-89, SHV-9, SHV-92, SHV-93, SHV-94, SHV-95, SHV-96, SHV-97, SHV-98, SHV-99 | (0.33, 1) | (0, 1) |

### D - SEQUENCE COMPARISON VERSUS ANNOTATION COMPARISON.

As explained in Section ”Materials and Methods”, to velidate Sarand, its extracted neighborhood sequences are compared against those of the reference genomes. However, the comparisons are based on matching Prokka gene annotations between underlying reference and extracted neighbourhoods. To explore whether annotation inconsistencies were leading to neighbourhoods being incorrectly labeled as false, we also directly compared the upstream/downstream neighborhood sequences. The sequence comparison was performed using BLASTN v2.9.0 and the threshold for identity and coverage was set to 90%. The comparison of these two cases for all simulated datasets in terms of precision and sensitivity are available in Figure 13 and Figure 14, respectively.

On the simple datasets 1_1_1 and 2_2_2, many more positive predictions were obtained by using sequence than when mapping annotations; the sensitivity on both datasets went from 76% and 34% to 90% respectively, while precision in both cases dropped below 50%. The difference on the CAMI_M datasets was never greater than 2%. The sequence-based approach yielded substantially higher precision (55% vs 92%) and sensitivity (67% vs 86%) on the CAMI_H_1 dataset. As an example, for a large group of SHV genes available in CAMI_H_1, Sarand can recover the sequence of the single neighborhood present in the reference fully with identity and coverage equal to 100 (i.e., precision = sensitivity = 1 for sequence comparison). Comparing annotations, no annotation was found for the upstream sequences extracted from the reference and Sarand (i.e., no neighborhood gene was found in the upstream sequence). However, regarding downstream sequences, given that Sarand’s sequence is longer than the reference sequence, it includes one extra gene which is not present in the reference annotation and makes Sarand’s annotation invalid (i.e., precision = sensitivity = 0 for annotation comparison). Supplementary Tables 3- 5 show different groups of AMR genes and Sarand’s performance when comparing annotations vs sequences.

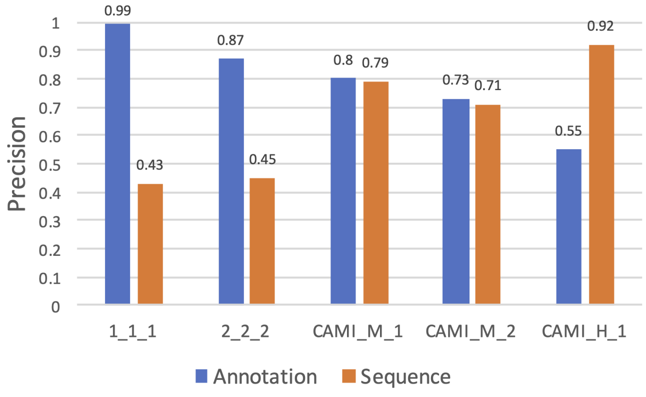


**Figure 13.** Precision of Sarand based on annotation vs sequence comparisons. For annotation evaluation, the gene-coverage threshold was set to 30, and for sequence evaluation the threshold for identity and coverage was set to 90%.

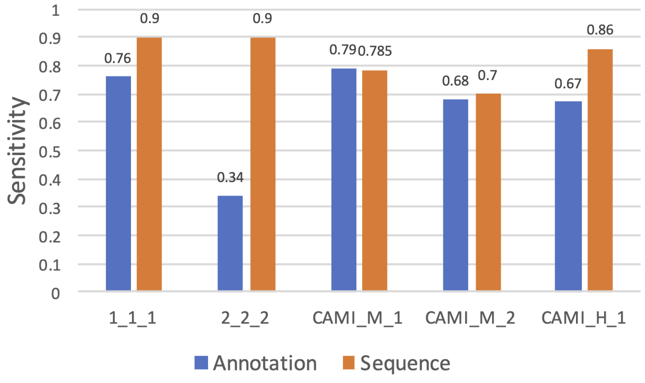


**Figure 14.** Sensitivity of Sarand based on annotation vs sequence comparisons. For annotation evaluation, the gene-coverage threshold was set to 30, and for sequence evaluation the threshold for identity and coverage was set to 90%.

**Table 3.** The difference between the performance results based on sequence comparison and annotation comparison for some AMR genes.

| Dataset | AMRs | Annotation | Sequence |
|---|---|---|---|
| CAMI_H_1 | SHV-42, SHV-43, SHV-44, SHV-45, SHV-46, SHV-48, SHV-49, SHV-5, SHV-50, SHV-51, SHV-52, SHV-53, SHV-55, SHV-56, SHV-57, SHV-59, SHV-6, SHV-60, SHV-61, SHV-62, SHV-63, SHV-64, SHV-65, SHV-66, SHV-67, SHV-69, SHV-7, SHV-70, SHV-71, SHV-72, SHV-73, SHV-74, SHV-105, SHV-128, SHV-151, SHV-18, SHV-41, SHV-75, SHV-129, SHV-13, SHV-133, SHV-134, SHV-135, SHV-137, SHV-14, SHV-140, SHV-141, SHV-142, SHV-143, SHV-144, SHV-145, SHV-147, SHV-148, SHV-149, SHV-15, SHV-150, SHV-76, SHV-77, SHV-78, SHV-79, SHV-8, SHV-80, SHV-81, SHV-82, SHV-83, SHV-84, SHV-85, SHV-86, SHV-89, SHV-9, SHV-92, SHV-93, SHV-94, SHV-95, SHV-96, SHV-97, SHV-98, SHV-99, SHV-180, SHV-182, SHV-183, SHV-185, SHV-186, SHV-187, SHV-188, SHV-189, SHV-19, SHV-2, SHV-20, SHV-21, SHV-22, SHV-23, SHV-24, SHV-25, SHV-26, SHV-27, SHV-28, SHV-29, SHV-2A, SHV-3, SHV-30, SHV-31, SHV-32, SHV-33, SHV-34, SHV-35, SHV-36, SHV-37, SHV-38, SHV-4, SHV-40, SHV-152, SHV-153, SHV-154, SHV-155, SHV-156, SHV-157, SHV-158, SHV-159, SHV-16, SHV-160, SHV-161, SHV-162, SHV-163, SHV-164, SHV-165, SHV-167, SHV-168, SHV-172, SHV-173, SHV-178, SHV-179, SHV-106, SHV-107, SHV-108, SHV-109, SHV-11, SHV-110, SHV-111, SHV-119, SHV-12, SHV-120, SHV-121, SHV-122, SHV-123, SHV-124, SHV-125, SHV-126, SHV-127, SHV-1, SHV-101, SHV-102, SHV-103, SHV-104 | $P=0, S=0$ | $P=1, S=1$ |
| Note: One neighborhood sequence has been extracted by Sarand. The sequence extracted from ref genomes matches sarand's sequence with identity=coverage=100. Regarding annotations, in upstream, no annotation was found in those two sequences. At downstream, Sarand's sequence is longer than reference sequence and includes one extra gene not available in reference annotation, which makes Sarand's sequence invalid. | | | |
| CAMI_H_1 | APH(3')-Ia | $P=0, S=0$ | $P=1, S=1$ |
| Note: two neighborhood sequences are extracted by sarand with the same upstream, while there are 2 sequences available in ref that one of them does not have anything in upstream. Comparing sequences, upstreams and downstreams match with identity and coverage above 99%. However, comparing annotations, with gene coverage $\text{threshold}=30$, genes are filtered by Sarand, and no match can be found. | | | |
| CAMI_H_1 | TEM-178, TEM-192, TEM-117, TEM-118, TEM-59, TEM-7, TEM-75, TEM-151, TEM-89, TEM-96, TEM-152, TEM-6, TEM-57, TEM-55, TEM-54, TEM-53, TEM-52, TEM-49, TEM-48, TEM-47, TEM-45, TEM-131, TEM-130, TEM-129, TEM-128, TEM-127, TEM-126, TEM-125, TEM-123, TEM-122, TEM-121, TEM-120, TEM-12, TEM-60, TEM-63, TEM-68, TEM-116, TEM-211, TEM-21, TEM-209, TEM-208, TEM-207, TEM-206, TEM-205, TEM-201, TEM-101, TEM-10, TEM-1, TEM-82, TEM-81, TEM-80, TEM-8, TEM-79, TEM-78, TEM-76, TEM-73, TEM-72, TEM-71, TEM-70, TEM-67, TEM-115, TEM-114, TEM-113, TEM-189, TEM-188, TEM-187, TEM-186, TEM-185, TEM-184, TEM-183, TEM-182, TEM-177, TEM-19, TEM-176, TEM-17, TEM-169, TEM-168, TEM-167, TEM-166, TEM-164, TEM-163, TEM-162, TEM-160, TEM-150, TEM-171, TEM-190, TEM-191, TEM-193, TEM-112, TEM-111, TEM-110, TEM-11, TEM-109, TEM-108, TEM-107, TEM-106, TEM-105, TEM-104, TEM-83, TEM-43, TEM-20, TEM-16, TEM-132, TEM-102, TEM-2, TEM-199, TEM-198, TEM-197, TEM-196, TEM-195, TEM-194, TEM-213, TEM-214, TEM-124, TEM-216, TEM-147, TEM-148, TEM-149, TEM-15, TEM-95, TEM-94, TEM-93, TEM-92, TEM-91, TEM-90, TEM-215, TEM-87, TEM-86, TEM-85, TEM-84, TEM-159, TEM-158, TEM-157, TEM-156, TEM-155, TEM-154, TEM-153, TEM-88, TEM-33, TEM-136, TEM-24, TEM-135, TEM-26, TEM-28, TEM-29, TEM-3, TEM-30, TEM-22, TEM-34, TEM-4, TEM-219, TEM-137, TEM-220, TEM-139, TEM-141, TEM-142, TEM-143, TEM-144, TEM-145, TEM-146, TEM-40, TEM-42, TEM-133, TEM-217, TEM-138, TEM-134 | $P=0.33, S=0.33$ | $P=1, S=0.75$ |
| Note: four neighborhood sequences were extracted by sarand (2 upstreams and 2 downstreams), and similarly 4 sequences were extracted from ref. Comparing sequences, both upstreams and both downstreams were valid, however out of 8 ref cases (4 up and 4 down seqs) only 6 cases were found by Sarand. Comparing annotations, only one of 3 found cases is valid and only one of 3 cases in ref was found. | | | |
| CAMI_H_1 | Escherichia_coli_ampC1_betalactamase, OXA-454, OXA-56, OXA-663, OXA-7, OXA-74, OXA-28, OXA-183, OXA-19, OXA-10, OXA-101, OXA-11, OXA-13, OXA-35, OXA-368, OXA-14, OXA-142, OXA-145, OXA-147, OXA-16, OXA-17, OXA-233, OXA-240, OXA-246, OXA-251, OXA-256 | $P=0.5, S=0.5$ | $P=0, S=0$ |
| Note: One neighborhood sequence has been extracted by sarand from the assembly graph and from the ref genomes. Comparing sequences, upstream sequences match with $\text{identity}=100$, $\text{coverage}=73$ and downstreams match with $\text{identity}=100$, $\text{coverage}=89$; therefore no true positive cases! Regarding annotations, downstreams are the same but in upstream, ref has one extra gene which is not available in Sarand (Sarand's upstream is shorter than ref upstream). | | | |
| CAMI_H_1 | QnrVC4, QnrVC5, QnrVC7 | $P=S=0$ | $P=0.33, S=0.5$ |
| CAMI_H_1 | tmrB | $P=S=0$ | $P=0.5, S=0.25$ |
| CAMI_H_1 | FosB | $P=S=0$ | $P=0.67, S=0.5$ |
| CAMI_H_1 | TEM-181 | $P=S=0$ | $P=S=0.33$ |
| CAMI_H_1 | sul2 | $P=S=0.33$ | $P=0.75, S=0.67$ |
| CAMI_H_1 | OXA-9, arnA | $P=S=1$ | $P=S=0.5$ |
| Note: One sequence has been extracted by sarand from the assembly graph and from the ref genomes. Comparing sequences, downstream sequences match perfectly but no hit can be found for upstream. Comparing annotations, no gene is annotated around the AMR in sequences of both sarand and ref; therefore $\text{precision}=\text{sensitivity}=1$. | | | |
| CAMI_H_1 | ANT(2")-Ia | $P=1, S=0.33$ | $P=0.5, S=0.25$ |
| CAMI_H_1 | SatA | $P=1, S=0.5$ | $P=0.5, S=0.5$ |
| CAMI_H_1 | aadA | $P=1, S=0.2$ | $P=0.75, S=0.57$ |

For annotation comparison, the gene-coverage threshold was set to 30, and for sequence comparison (using Blastn) the identity and coverage thresholds were set to 90%. P and S refer to precision and sensitivity, respectively.

**Table 4.** The difference between the performance results based on sequence comparison and annotation comparison for some AMR genes.

| Dataset | AMRs | Annotation | Sequence |
|---|---|---|---|
| 1_1_1 | TEM-104, TEM-105, TEM-106, TEM-107, TEM-108, TEM-109, TEM-11, TEM-110, TEM-111, TEM-112, TEM-113, TEM-114, TEM-115, TEM-116, TEM-12, TEM-120, TEM-121, TEM-122, TEM-123, TEM-124, TEM-125, TEM-199, TEM-2, TEM-20, TEM-201, TEM-205, TEM-206, TEM-207, TEM-208, TEM-209, TEM-21, TEM-211, TEM-213, TEM-214, TEM-215, TEM-216, TEM-217, TEM-219, TEM-22, TEM-220, TEM-102, TEM-126, TEM-148, TEM-17, TEM-198, TEM-24, TEM-55, TEM-149, TEM-15, TEM-150, TEM-151, TEM-152, TEM-153, TEM-154, TEM-155, TEM-156, TEM-157, TEM-158, TEM-159, TEM-16, TEM-160, TEM-162, TEM-163, TEM-164, TEM-166, TEM-167, TEM-168, TEM-169, TEM-57, TEM-59, TEM-6, TEM-60, TEM-63, TEM-67, TEM-68, TEM-70, TEM-71, TEM-72, TEM-73, TEM-76, TEM-78, TEM-79, TEM-8, TEM-80, TEM-81, TEM-82, TEM-83, TEM-84, TEM-85, TEM-86, TEM-87, TEM-88, TEM-90, TEM-91, TEM-92, TEM-93, TEM-94, TEM-95, TEM-96, TEM-1, TEM-10, TEM-101, TEM-127, TEM-128, TEM-129, TEM-130, TEM-131, TEM-132, TEM-133, TEM-134, TEM-135, TEM-136, TEM-137, TEM-138, TEM-139, TEM-141, TEM-142, TEM-143, TEM-144, TEM-145, TEM-146, TEM-147, TEM-171, TEM-176, TEM-177, TEM-178, TEM-182, TEM-183, TEM-184, TEM-185, TEM-186, TEM-187, TEM-188, TEM-189, TEM-19, TEM-190, TEM-191, TEM-193, TEM-194, TEM-195, TEM-196, TEM-197, TEM-26, TEM-28, TEM-29, TEM-3, TEM-30, TEM-33, TEM-34, TEM-4, TEM-40, TEM-42, TEM-43, TEM-45, TEM-47, TEM-48, TEM-49, TEM-52, TEM-53, TEM-54, TEM-217 | $P=1$, $S=0.8$ | $P=0.24$, $S=1$ |
| 1_1_1 | SHV-183, SHV-185, SHV-186, SHV-187, SHV-188, SHV-189, SHV-19, SHV-2, SHV-20, SHV-21, SHV-22, SHV-23, SHV-24, SHV-25, SHV-26, SHV-27, SHV-28, SHV-29, SHV-2A, SHV-3, SHV-30, SHV-31, SHV-32, SHV-33, SHV-34, SHV-35, SHV-108, SHV-147, SHV-182, SHV-36, SHV-61, SHV-81, SHV-62, SHV-63, SHV-64, SHV-65, SHV-66, SHV-67, SHV-69, SHV-7, SHV-70, SHV-71, SHV-72, SHV-73, SHV-74, SHV-75, SHV-76, SHV-77, SHV-78, SHV-79, SHV-8, SHV-80, SHV-109, SHV-11, SHV-110, SHV-111, SHV-119, SHV-12, SHV-120, SHV-121, SHV-122, SHV-123, SHV-124, SHV-125, SHV-126, SHV-127, SHV-128, SHV-129, SHV-13, SHV-133, SHV-134, SHV-135, SHV-137, SHV-14, SHV-140, SHV-141, SHV-142, SHV-143, SHV-144, SHV-145, SHV-1, SHV-100, SHV-101, SHV-102, SHV-103, SHV-104, SHV-105, SHV-106, SHV-107, SHV-148, SHV-149, SHV-15, SHV-150, SHV-151, SHV-152, SHV-153, SHV-154, SHV-155, SHV-156, SHV-157, SHV-158, SHV-159, SHV-16, SHV-160, SHV-161, SHV-162, SHV-163, SHV-164, SHV-165, SHV-167, SHV-168, SHV-172, SHV-173, SHV-178, SHV-179, SHV-18, SHV-180, SHV-37, SHV-38, SHV-4, SHV-40, SHV-41, SHV-42, SHV-43, SHV-44, SHV-45, SHV-46, SHV-48, SHV-49, SHV-5, SHV-50, SHV-51, SHV-52, SHV-53, SHV-55, SHV-56, SHV-57, SHV-59, SHV-6, SHV-60, SHV-82, SHV-83, SHV-84, SHV-85, SHV-86, SHV-89, SHV-9, SHV-92, SHV-93, SHV-94, SHV-95, SHV-96, SHV-97, SHV-98, SHV-99 | $P=1$, $S=0.67$ | $P=0.5, S=0.75$ |
| 1_1_1 | dfrA14 | $P=S=0$ | $P=0.17$, $S=0.5$ |
| 1_1_1 | Staphylococcus_aureus_FosB | $P=S=0.5$ | $P=0.67$, $S=1$ |
| 1_1_1 | APH(3')-Ia | $P=1$, $S=0.33$ | $P=0.27$, $S=1$ |
| 1_1_1 | aadA12, aadA17, aadA2, aadA25, aadA3, aadA8, aadA8b | $P=1$, $S=0.5$ | $P=0.5, S=1$ |
| 1_1_1 | catII_from_Escherichia_coli_K-12, catI, plasmid-encoded_cat_(pp-cat) | $P=1$, $S=0.5$ | $P=0.6, S=1$ |
| 1_1_1 | AAC(6')-Ie-APH(2")-Ia | $P=1$, $S=0.5$ | $P=1, S=1$ |
| 1_1_1 | TEM-117 | $P=S=0.5$ | $P=0.24$, $S=1$ |
| 2_2_2 | TEM-185, TEM-155, TEM-154, TEM-153, TEM-152, TEM-151, TEM-150, TEM-149, TEM-156, TEM-145, TEM-139, TEM-138, TEM-137, TEM-134, TEM-133, TEM-131, TEM-130, TEM-142, TEM-129, TEM-157, TEM-16, TEM-8, TEM-78, TEM-3, TEM-26, TEM-24, TEM-22, TEM-219, TEM-159, TEM-213, TEM-21, TEM-205, TEM-20, TEM-168, TEM-167, TEM-162, TEM-160, TEM-211, TEM-80, TEM-125, TEM-121, TEM-17, TEM-146, TEM-120, TEM-199, TEM-197, TEM-195, TEM-194, TEM-2, TEM-193, TEM-190, TEM-19, TEM-189, TEM-188, TEM-187, TEM-184, TEM-183, TEM-191, TEM-123, TEM-73, TEM-40, TEM-72, TEM-71, TEM-68, TEM-67, TEM-63, TEM-60, TEM-6, TEM-4, TEM-59, TEM-53, TEM-52, TEM-49, TEM-48, TEM-47, TEM-45, TEM-42, TEM-54, TEM-178, TEM-81, TEM-83, TEM-82, TEM-171, TEM-110, TEM-11, TEM-109, TEM-101, TEM-1, TEM-96, TEM-94, TEM-93, TEM-91, TEM-90, TEM-89, TEM-87, TEM-86, TEM-85, TEM-84, TEM-92, TEM-112, TEM-111, TEM-114, TEM-113, TEM-12, TEM-116, TEM-115, TEM-177 | $P=1$, $S=0.08$ | $P=0.27$, $S=0.93$ |
| 2_2_2 | CTX-M-126, CTX-M-125, CTX-M-122, CTX-M-121, CTX-M-113, CTX-M-112, CTX-M-111, CTX-M-110, CTX-M-106, CTX-M-105, CTX-M-104, CTX-M-102, CTX-M-87, CTX-M-86, CTX-M-85, CTX-M-129, CTX-M-24, CTX-M-130, CTX-M-99, CTX-M-98, CTX-M-93, CTX-M-90, CTX-M-21, CTX-M-19, CTX-M-17, CTX-M-161, CTX-M-16, CTX-M-159, CTX-M-148, CTX-M-147, CTX-M-14, CTX-M-134, CTX-M-13, CTX-M-83, CTX-M-84, CTX-M-81, CTX-M-67, CTX-M-65, CTX-M-51, CTX-M-50, CTX-M-49, CTX-M-48, CTX-M-46, CTX-M-47, CTX-M-38, CTX-M-27, CTX-M-9, CTX-M-45 | $P=0, S=0$ | $P=0.67$, $S=1$ |
| 2_2_2 | TEM-176, TEM-182, TEM-186, TEM-196, TEM-198, TEM-33, TEM-34, TEM-43, TEM-55, TEM-57, TEM-70, TEM-122, TEM-124, TEM-126, TEM-127, TEM-128, TEM-132, TEM-135, TEM-136, TEM-141, TEM-143, TEM-144, TEM-147, TEM-148, TEM-15, TEM-158, TEM-163, TEM-164, TEM-166, TEM-169, TEM-201, TEM-206, TEM-207, TEM-208, TEM-209, TEM-214, TEM-215, TEM-216, TEM-217, TEM-220, TEM-28, TEM-29, TEM-30, TEM-76, TEM-79, TEM-88, TEM-95, TEM-10, TEM-102, TEM-104, TEM-105, TEM-106, TEM-107, TEM-108, TEM-192 | $P=1$, $S=0.09$ | $P=0.27$, $S=0.93$ |
| 2_2_2 | AAC(6')-Ib-cr, AAC(6')-Ib3, AAC(6')-Ib11, AAC(6')-Ib', AAC(6')-Ib10, AAC(6')-Ib, AAC(6')-Ib-Suzhou, AAC(6')-Ib-Hangzhou, AAC(6')-Ib4, AAC(6')-Ib9 | $P=0.5, S=0.5$ | $P=0.33$, $S=1$ |
| 2_2_2 | KPC-11, KPC-12, KPC-13, KPC-14, KPC-15, KPC-16, KPC-17, KPC-19, KPC-22, KPC-24, KPC-3, KPC-4, KPC-5, KPC-6, KPC-7, KPC-8, KPC-9, KPC-1 | $P=1, S=1$ | $P=0.33$, $S=1$ |
| 2_2_2 | dfrA14 | $P=S=0$ | $P=0.25$, $S=0.5$ |

For annotation comparison, the gene-coverage threshold was set to 30, and for sequence comparison (using Blastn) the identity and coverage thresholds were set to 90%. P and S refer to precision and sensitivity, respectively.

**Table 5.** The difference between the performance results based on sequence comparison and annotation comparison for some AMR genes.

| Dataset | AMRs | Annotation | Sequence |
|---|---|---|---|
| 2_2_2 | TEM-181 | $P=1$, $S=0.08$ | $P=0.3, S=0.86$ |
| 2_2_2 | TEM-7, TEM-75, TEM-117, TEM-118 | $P=1$, $S=0.1$ | $P=0.27$, $S=0.93$ |
| 2_2_2 | sul1 | $P=0.14$, $S=0.33$ | $P=0.38$, $S=0.75$ |
| 2_2_2 | OXA-9 | $P=S=0.5$ | $P=0.29$, $S=1$ |
| 2_2_2 | Staphylococcus_aureus_FosB | $P=S=0.5$ | $P=0.67$, $S=1$ |
| 2_2_2 | APH(3')-Ia | $P=1$, $S=0.33$ | $P=0.08$, $S=0.25$ |
| 2_2_2 | catI, plasmid-encoded_cat_(pp-cat) | $P=1$, $S=0.25$ | $P=0.22$, $S=0.5$ |
| 2_2_2 | catII_from_Escherichia_coli_K-12 | $P=1$, $S=0.5$ | $P=0.25$, $S=0.5$ |
| CAMI_M_1 | lsaB | $P=S=0$ | $P=S=0.5$ |
| CAMI_M_1 | AAC(6')-Ii | $P=S=1$ | $P=S=0$ |
| Note: One neighborhood sequence has been extracted by sarand from the assembly graph and from the ref genomes. Comparing sequences, downstreams match with $\mathrm{identity}=99$, $\mathrm{coverage}=87$; so no true positive case! comparing annotations, no gene around AMR was found so $\mathrm{precision}=\mathrm{sensitivity}=1$. | | | |
| CAMI_M_2 | lsaB | $P=S=1$ | $P=S=0.5$ |
| CAMI_M_2 | ugd | $P=1$, $S=0.5$ | $P=0.33$, $S=0.5$ |
| CAMI_M_2 | Escherichia_coli_ampC1_beta-lactamase | $P=S=0.5$ | $P=S=1$ |

For annotation comparison, the gene-coverage threshold was set to 30, and for sequence comparison (using Blastn) the identity and coverage thresholds were set to 90%. P and S refer to precision and sensitivity, respectively.
